# A Generalized Spatial Progress Code for Navigation in the Medial Prefrontal Cortex

**DOI:** 10.64898/2026.07.02.735049

**Authors:** Yizheng Wang, Lingzhu Fan, Min Xie, Danhua Wan, Shanshan Liu, Feifan Sun, Jun Tian, Baorong Zhang, Lixia Gao, Xinjian Li

**Author notes:** These authors contributed equally (Yizheng Wang, Lingzhu Fan, Min Xie).

## Abstract

Goal-directed navigation requires an internal estimate of progress toward a desired outcome, yet how the brain constructs and represents such a signal remains unclear. Here, we designed a multi-route navigation paradigm with geometrically distinct paths and imaged medial prefrontal cortex (mPFC) activity as mice performed goal-directed route choice behavior. We found that a substantial fraction of mPFC neurons fired at consistent positions along a normalized start-to-goal axis, independent of path geometry, running direction, and task context. At the ensemble level, mPFC activity reliably decoded spatial progress but carried minimal information about path identity. This progress code emerged with task familiarity and required input from the hippocampal CA1. Inactivation of either mPFC or CA1→mPFC projection did not disrupt learned path preference but impaired navigational fluency in trained mice. These findings reveal a hippocampal-prefrontal mechanism for representing abstract, context-generalized spatial progress within a goal-directed navigational episode, supporting fluent navigation across variable routes.

## INTRODUCTION

Efficient goal-directed spatial navigation in natural environments requires individuals to recognize allocentric space, choose optimal paths and estimate relative progress from start to goal. Previous studies have identified the hippocampus, entorhinal cortex and related brain structures as core regions to build an internal allocentric cognitive map(Behrens et al., 2018; Buzsáki and Moser, 2013; Epstein et al., 2017; O’Keefe and Dostrovsky, 1971; Schiller et al., 2015). The concerted activity of place cells, grid cells, border cells and corner cells within this circuit dynamically represents an individual’s precise location(Behrens et al., 2018; Fyhn et al., 2004; Moser et al., 2008; Moser et al., 2017; O’Keefe and Dostrovsky, 1971; Solstad et al., 2008; Sun et al., 2024). The retrosplenial cortex and parahippocampal cortex have also been recognized as essential for scene recognition and landmark anchoring(Epstein, 2008; Yoder et al., 2011). In addition, several brain regions, including the medial prefrontal cortex (mPFC), the hippocampus (HPC), and the secondary motor cortex have been proposed to function in route choice, as evidenced by biased neural activity during two-way route selection(Ito et al., 2015; Muysers et al., 2025; Olson et al., 2020). However, efficient navigation in natural environments with multiple paths also requires individuals to estimate relative progress from start to goal, raising an unsolved issue: “How much have I accomplished?”

The mPFC is an important high-level cognitive region critical for spatial navigation. Previous studies have revealed that mPFC receives anatomical projections from the hippocampus(Euston et al., 2012; Ge et al., 2023; Jay and Witter, 1991; Qiu et al., 2024; Sauer et al., 2022; Spellman et al., 2015; Tavares and Tort, 2022) and that its disruption impairs spatial navigation in both rodents(de Bruin et al., 1994; Granon and Poucet, 1995) and humans(Ciaramelli, 2008). In addition, mPFC neurons exhibit diverse representations of task space, including current spatial location(Ma et al., 2023; Zielinski et al., 2019), goal locations(Hok et al., 2005; Vogel et al., 2022), specific trajectory routes(Fujisawa et al., 2008; Ito et al., 2015), task sequences(Tang et al., 2023), and value-based decision making(Veselic et al., 2025). However, it remains unclear how mPFC contributes to route choice behavior in naturalistic space with multiple paths and whether it estimates the relative navigational progress.

Here, we designed a route choice paradigm that incorporates multiple geometrically distinct paths to mimic real-world spatial navigation. By imaging mPFC neural activity when mice were freely navigating a multi-path maze, we addressed: (1) whether mPFC is involved in route choice behavior in multi-path space; (2) whether mPFC estimates the relative navigational progress; and (3) the origin of the progress representation. Our results reveal that the mPFC provides a spatial-progress scaffold: not a map of physical space, and not a simple path selector, but an abstract representation of goal-directed advancement, illustrating how the brain integrates diverse spatial experiences into a unified internal metric of progress for flexible and efficient spatial navigation.

## RESULTS

### The mPFC supports navigational fluency but not path preference of learned route choice behavior

To investigate route choice behavior in mice within a naturalistic and complex spatial environment, we established a route choice paradigm equipped with two water ports and four geometrically different paths: a long curve, a zigzag path, a direct path, and a short curve (Figures 1A and S1A). Mice were motivated to select different paths and navigate back and forth between the two water ports. Different paths represent distinct travel strategies: for instance, the direct path provides the shortest and straightest trajectory, whereas the long curve is the longest and most circuitous. The whole task lasted 15 sessions (15 min/session, 1 session/day) in which the proportion of incomplete trials and trial duration decreased gradually (Figures S1B and S1C). Route choice behavior showed two relatively stable stages during training, which were defined as the naive (sessions 3–5) and trained (sessions 13–15) stages (Figures 1A and S1B–S1D). Compared with the naive stage, mice in the trained stage completed more trials with a lower proportion of incomplete trials, shorter trial duration, and reduced choice time in the route choice regions in a session (Figures 1B–1E and S1E–S1G). Trained mice also displayed a lower proportion of random pause, indicating enhanced task execution stability (Figures 1F, S1H, and S1I). Importantly, whereas route choice was uniformly distributed during the naive stage, trained mice exhibited a significant bias toward the direct path while eschewing the long curve (Figures 1G and 1H), resulting in a higher path preference index (Figure 1I). Together, these results indicate that training-induced task familiarity optimizes route choice behavior, supporting efficient goal-directed spatial navigation.

**Figure 1.**
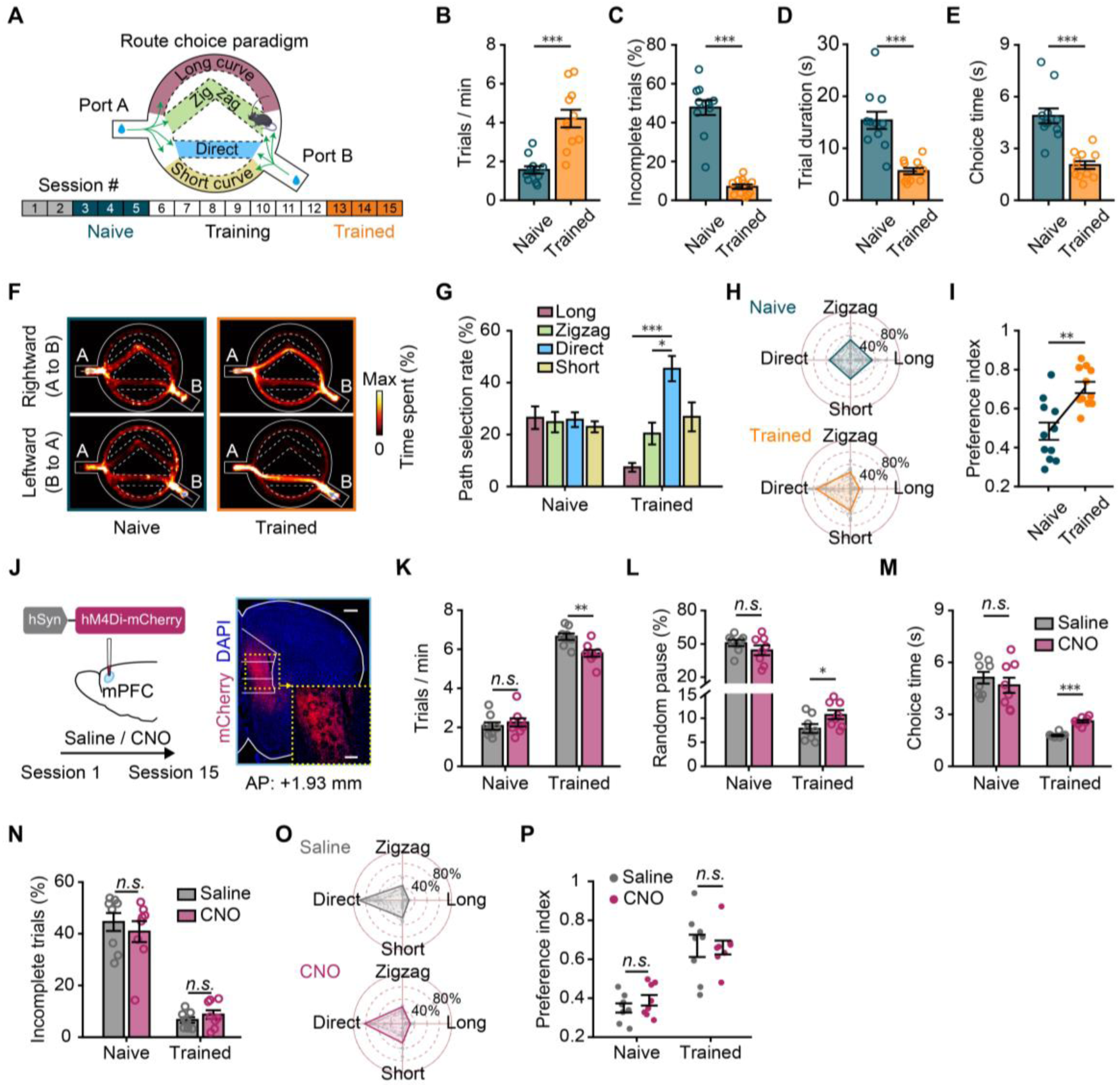
The mPFC supports navigational fluency but not path preference in trained mice. (A) Schematic showing the route choice paradigm (top) and training timeline (bottom). (B–E) Comparison of trials per minute (B), proportion of incomplete trials (C), trial duration (D), and route choice time (E) between the naive and trained stages (*n* = 11 mice) (F) Representative heatmaps showing the movement of mice in the naive and trained stages. Top, rightward trials; bottom, leftward trials. (G–I) Comparison of path selection rate (G, H) and preference index (I) between the naive and trained stages (*n* = 11 mice). (J) Left, schematic showing the mPFC inactivation strategy. Saline or CNO administration was performed before each session throughout the experiment. Right, representative fluorescence image showing expression of hM4Di-mCherry in mPFC. Scale bars: main: 500 µm, inset: 200 µm. (K–N) Comparison of trials per minute (K), proportion of random pause (L), route choice time (M), and proportion of incomplete trials (N) between the saline and CNO groups (*n* = 8 mice per group). (O, P) Comparison of path selection rate (O, the naive stage is shown in Figure S1I) and preference index (P) between the saline and CNO groups (*n* = 8 mice per group). Values are expressed as mean ± s.e.m. For statistical significance, \**P*<0.05, \*\**P*<0.01 and \*\*\**P*<0.001. *n.s.*, not significant. Detailed statistics are presented in Table S1. See also Figure S1.

To examine the role of the mPFC in route choice behavior, we bilaterally expressed the inhibitory DREADD human Gi-coupled M4 muscarinic receptor (hM4Di-mCherry) in the mPFC and administered the clozapine N-oxide (CNO; 10 mg/kg, i.p.) before each task session to inactivate mPFC activity (Figure 1J). While no difference was observed between the saline and CNO groups in the naive stage, mice treated with CNO completed fewer trials, exhibited a higher proportion of random pause and longer route choice time in the trained stage (Figures 1K–1M). Notably, the proportion of incomplete trials, path preference, and inter-trial interval remained unaffected after inactivation of the mPFC (Figures 1N–1P, S1J, and S1K). Thus, the mPFC primarily facilitates fluent navigation rather than determining detailed path selection.

### Encoding of spatial progress in the mPFC

To decipher how the mPFC encodes spatial navigation in a complex multi-path environment, we expressed the fluorescent calcium indicator GCaMP6s in mPFC excitatory neurons and imaged their activity using a miniature fluorescence microscope in both naive and trained stages (Figures 2A and S2A–S2F; Video S1). Notably, a subset of neurons was found to maintain a stable and sequential firing pattern across consecutive trials, regardless of the running direction or the chosen path (Figure 2B; Video S2). To compare neural activity across trajectories with different geometries, we linearized each trial into normalized spatial-progress bins, thereby registering mPFC activity by relative spatial progress rather than absolute spatial position (Figures S2G and S2H; STAR Methods). We next compared the progress-aligned response profiles between two running directions and found a clear activation band of the leftward trials in the trained stage, which resulted in a clear diagonal correlation coefficient band that was absent in the naive stage (Figures S3A and S3B). The correlation coefficients of neuronal activity between two running directions were significantly higher after training (Figure S3C) and exceeded those obtained after reversing or shuffling the activity in the leftward trials (Figure S3D). Similar increases were also observed across different paths and across repeated trials of the same path type both within and across days (Figures S3E and S3F). Together, these results indicate that training drives the emergence of a progress-aligned representation in the mPFC that generalizes across running directions and geometrically distinct paths.

**Figure 2.**
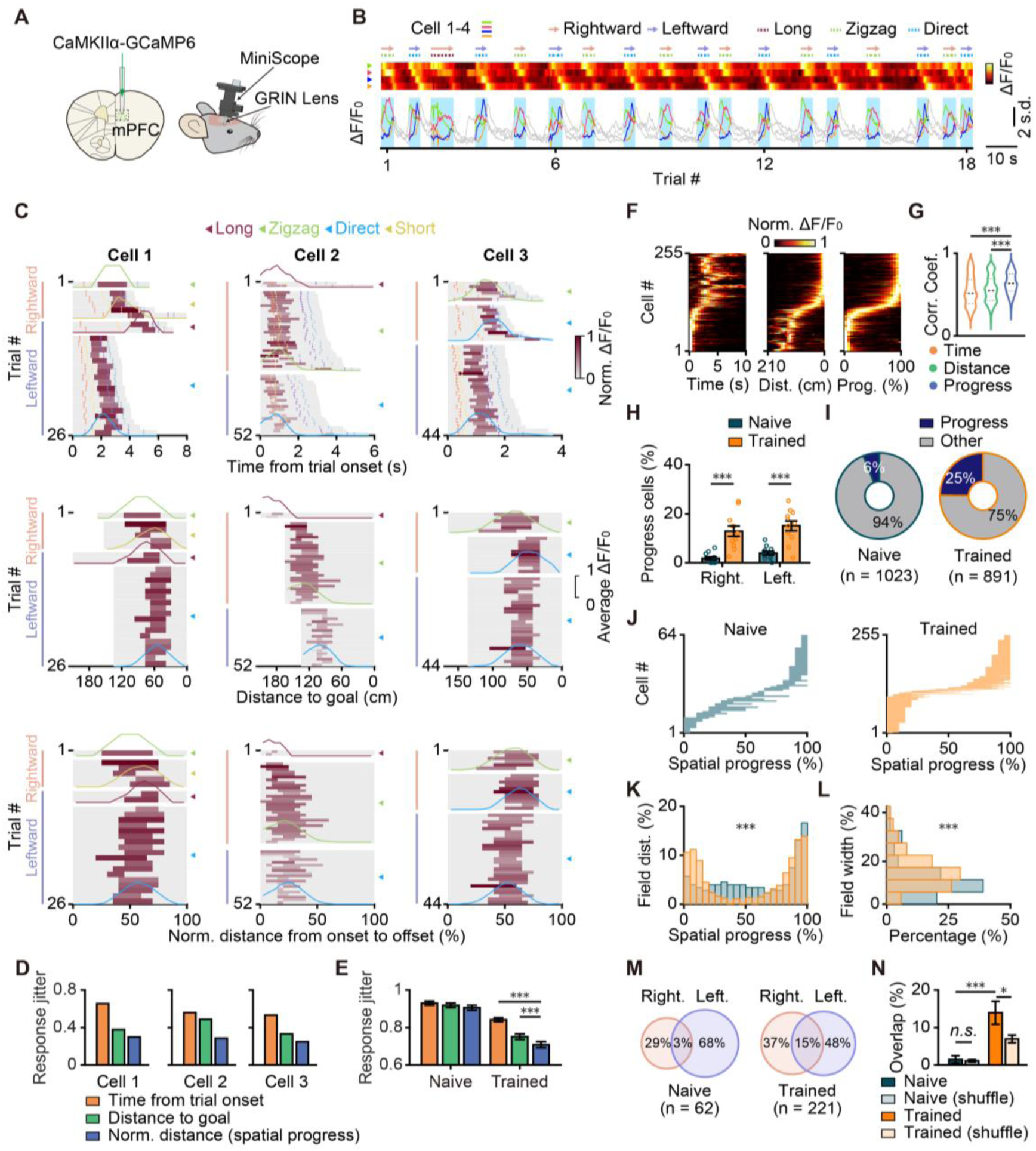
The mPFC encodes spatial progress at the single-cell level. (A) Schematic showing the virus injection plan (left) and imaging setup (right). (B) Activity profiles of four representative mPFC neurons sorted by their mean peak firing time across trials. Top, heatmap. Bottom, calcium traces. Arrows and dashed lines at the top indicate the running direction and chosen path, respectively. Vertical light blue shadows denote individual trial epochs. (C) Activity maps of three example mPFC neurons as a function of time from trial onset (top), distance to goal (middle), or normalized distance from trial onset to offset (bottom). Overlaid curves show the average activity calculated separately from trials of each path type. Colored dashed lines mark behavioral events: leave reward (orange), path start (yellow), path end (purple), and enter reward (blue). (D) Comparison of the response jitter across different alignment conditions for the example neurons in (C). (E) Comparison of the response jitter for the recorded mPFC neurons across different alignment conditions between the naive and trained stages (naive, *n* = 1023 cells; trained, *n* = 891 cells). (F) Trial-averaged activity maps of the identified spatial-progress cells aligned by time from trial onset (left), distance to goal (middle), or spatial progress (right). Cells in all three conditions are sorted by peak position of progress-aligned activity. Cells classified as spatial-progress cells for both directions are shown twice. (G) Correlation coefficients calculated by correlating each neuron’s activity in individual trials with its trial-averaged activity (*n* = 255 spatial-progress cells in the trained stage). (H) Proportions of spatial-progress cells for rightward and leftward trials in the naive and trained stages (naive, *n* = 1023 cells; trained, *n* = 891 cells). (I) Proportions of overall spatial-progress cells. (J–L) Progress fields (J), field distributions (K), and field widths (L) of spatial-progress cells in the naive and trained stages (naive, *n* = 64 cells, trained, *n* = 255 cells). (M) Venn diagrams illustrating the relationship of spatial-progress cells between rightward and leftward trials in the naive and trained stages. (N) Comparison of the proportion of overlapped spatial-progress cells (*n* = 11 mice). \**P*<0.05, \*\**P*<0.01 and \*\*\**P*<0.001. *n.s.*, not significant.Detailed statistics are presented in Table S1. See also Figures S2–S5 and Video S1.

The highly consistent neuronal activity across different paths led us to hypothesize that the mPFC encodes spatial progress, distinct from elapsed time or distance to goal. To test this, we aligned the activity dynamics of individual neurons according to three variables: elapsed time from trial onset, distance to goal, or spatial progress (normalized distance from onset to offset) (Figures 2C and S4). Alignment by elapsed time caused the responses to shift toward trial offset as trial duration increased, whereas alignment by distance to goal caused them to shift toward trial onset as path length increased. By contrast, alignment by spatial progress resulted in similar relative peak positions across path geometries and running directions (Figures 2C and S4). We next quantified response consistency across the three alignment conditions using a jitter-based analysis (STAR Methods). In the trained, but not naive, stage, jitter was lowest when activity was aligned by spatial progress compared with the other two conditions (Figures 2D and 2E). Thus, the activity patterns of these neurons were better explained by a representation of spatial progress from start to goal, rather than by elapsed time or distance to goal. We therefore termed these neurons spatial-progress cells.

To identify the spatial-progress cells, we selected neurons that showed place-cell-like tuning within individual paths and peaked at similar normalized positions across geometrically distinct paths (Figure S5A; STAR Methods). Using these criteria, we identified a large proportion of spatial-progress cells in the trained stage. Compared with alignment by elapsed time or distance to goal, alignment by spatial progress produced narrower response fields and higher correlations between single-trial activity and trial-averaged activity (Figures 2F and 2G), indicating that spatial progress best accounted for the activity of these neurons.

Training significantly increased the proportion of spatial-progress cells in both running directions, comprising approximately 25% of the recorded population in the trained stage (Figures 2H and 2I). These neurons showed no significant spatial clustering within the imaging field of view (Figures S5B and S5C). Although their progress fields were primarily clustered near trial offset in the naive stage and became enriched near both trial onset and offset in the trained stage, they spanned the full navigational trajectory (Figures 2J and 2K). Progress-field width was approximately 15% of the normalized path in the trained stage and was significantly greater than in the naive stage (Figure 2L). In addition, the proportion of spatial-progress cells active in both running directions increased after training and significantly exceeded chance levels, whereas no significant bidirectional overlap was observed in the naive stage (Figures 2M and 2N). Together, these results demonstrate that the mPFC encodes spatial progress at the single-cell level, and that training increases both the prevalence and direction-generalization of spatial-progress cells.

### The mPFC population activity decodes route-invariant spatial progress

Given the spatial-progress coding at the single-cell level, we next asked whether the mPFC population activity was organized by spatial progress. We binned neuronal activity according to spatial progress and visualized the resulting activity patterns using the dimensionality reduction technique uniform manifold approximation and projection (UMAP) separately for each running direction (Figures 3A and S6; STAR Methods). Activity patterns from similar progress stages clustered together, whereas those from different stages were more separated, as reflected by shorter Euclidean distances within the same progress stage than between different stages (Figure 3E). We then connected activity patterns across consecutive progress bins to construct single-trial neural trajectories, from which the path-averaged trajectories were derived (Figures 3B and S7). Neural trajectories from different trials and paths followed a similar ring-like structure, with matched progress stages closely aligned across paths. Consistently, Euclidean distances did not differ between same-path and different-path trajectories at matched progress stages (Figure 3F). The separation between successive progress bins was greatest near trial onset and offset (Figure 3C), consistent with the enrichment of progress fields at these phases (Figures 2J and 2K). When trajectories from both running directions were analyzed together, activity patterns carried the directional information, as reflected by greater separation between same-direction and opposite-direction trajectories (Figures 3D and 3G). These structured trajectories were absent in the naive stage (Figures S6 and S7) and remained consistent across consecutive trials even when mice switched direction or path (Figure S8; Video S2). Together, these results indicate that mPFC activity follows a reproducible spatial-progress trajectory after training that is largely invariant to path geometry.

**Figure 3.**
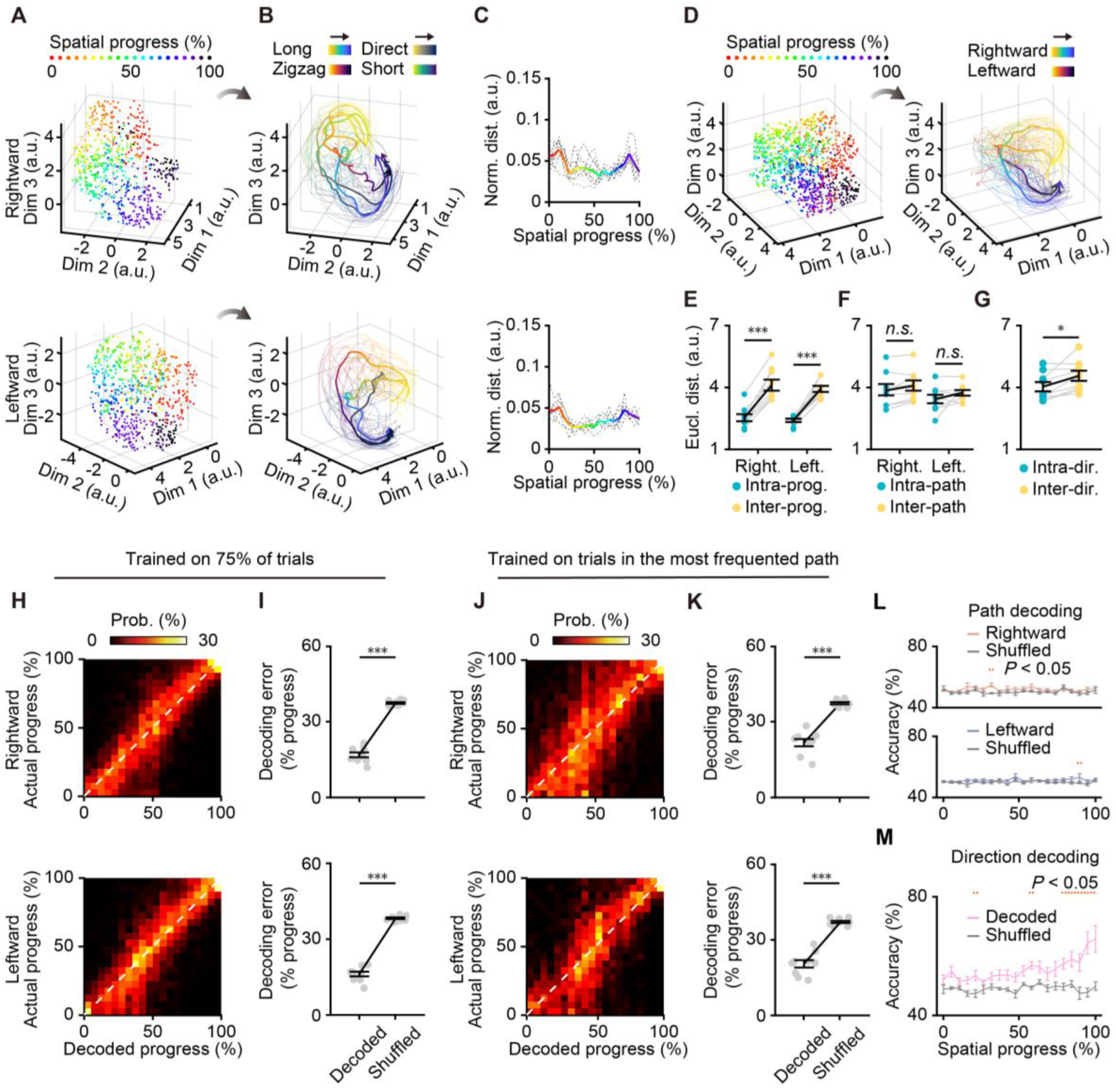
mPFC activity decodes the spatial progress instead of the selected path. (A) Neuronal activity (*n* = 195 cells) from an example animal binned into spatial-progress bins for trials of different paths and visualized on a neural manifold (UMAP). Manifolds were constructed from rightward (top) and leftward (bottom) trials. Colored dots denote the spatial progress. (B) Neural trajectories on the manifolds constructed from (A), grouped by paths and color-coded by both path type and spatial progress. Thick arrows, direction of the mean trajectory for each path along spatial progress. Fine lines, neural trajectories of individual trials. (C) Normalized distance between successive bins along the trajectories in (B), color-coded by spatial progress. Each dashed gray line represents one animal. (D) Left, manifolds constructed from all trials. Right, trajectories grouped by directions.\ (E) Average Euclidean distance within and between spatial-progress stages (*n* = 9 mice). (F) Average Euclidean distance between trajectory pairs of the same versus different paths (*n* = 9 mice). (G) Average Euclidean distance between trajectory pairs of the same versus different running directions (*n* = 9 mice). (H) Confusion matrices showing the decoded versus actual spatial progress using 75% of randomly chosen trials as the training dataset. Top, rightward trials; bottom, leftward trials. (I) Comparison of the decoding error between actual and shuffled datasets from (H) (*n* = 9 mice). (J) Confusion matrices showing the decoded versus actual spatial progress using trials from the most frequented path as the training dataset and trials from the second-most frequented path as the test dataset. Top, rightward trials; bottom, leftward trials. (K) Comparison of the decoding error between actual and shuffled datasets from (J) (*n* = 9 mice). (L) Decoding accuracy for path types using neuronal activity from individual progress bins. The performance was compared bin by bin with the chance level (*n* = 9 mice). Dashed lines at the top indicate bins with decoding accuracy significantly higher than chance level. (M) Same as (L), but running directions were decoded (*n* = 9 mice). \**P*<0.05, \*\**P*<0.01 and \*\*\**P*<0.001. *n.s.*, not significant. Detailed statistics are presented in Table S1. See also Figures S6–S9 and Video S2.

To test whether spatial progress could be reliably decoded from mPFC activity, we trained linear support vector machine (SVM) models using binned population activity from 75% of randomly selected trials in each direction and predicted the spatial progress using the remaining trials (STAR Methods). Decoded progress closely approximated the actual progress, producing clear diagonal bands in the confusion matrices and significantly lower decoding error than shuffled controls for both running directions (Figures 3H and 3I). In addition, decoding performance remained above chance whether models were trained on the most frequented path and tested on the second-most frequented path, trained and tested on different halves of trials, or restricted to 10%–90% spatial progress (Figures 3J, 3K, and S9). These results show that mPFC activity reliably encodes spatial progress across trials and paths. Finally, using a naive Bayes classifier, we asked whether mPFC activity encoded path identity or running direction (STAR Methods). Path identity could not be decoded above chance at any progress stage in either direction (Figure 3L). In contrast, running direction was decoded above chance throughout the task (Figure 3M). Thus, mPFC activity primarily represents spatial progress and running direction, rather than the specific path chosen. Overall, these findings demonstrate that mPFC activity encodes a sequential, route-invariant representation of spatial progress.

### The mPFC spatial-progress coding generalizes across routes and contexts

In our paradigm, mice exhibited a strong path preference in the trained stage, with relatively few choices of the long curve, raising the possibility that uneven path sampling could limit the assessment of spatial-progress generalization. To address this, we modified the paradigm into a no-choice task, in which only one path was accessible per session and the open path alternated across four consecutive sessions (Figure 4A). In this setting, the normalized peak positions of individual neurons remained aligned across paths and differed from shuffled controls (Figure 4B). Correlations of peak positions and binned neuronal activity were significantly higher across path pairs and running directions than in shuffled datasets (Figures 4C, 4D and S10A–S10C). Using the same criteria as in the free-choice task, we found that approximately 16% of neurons showed consistent spatial-progress coding across paths and sessions (Figures 4E, 4F, S10D, and S10E). Moreover, SVM models trained on trials from three paths accurately decoded spatial progress in the remaining path, with significantly lower decoding error than shuffled controls (Figure 4G). These results indicate that mPFC spatial-progress coding generalizes across paths even when path availability is experimentally constrained.

**Figure 4.**
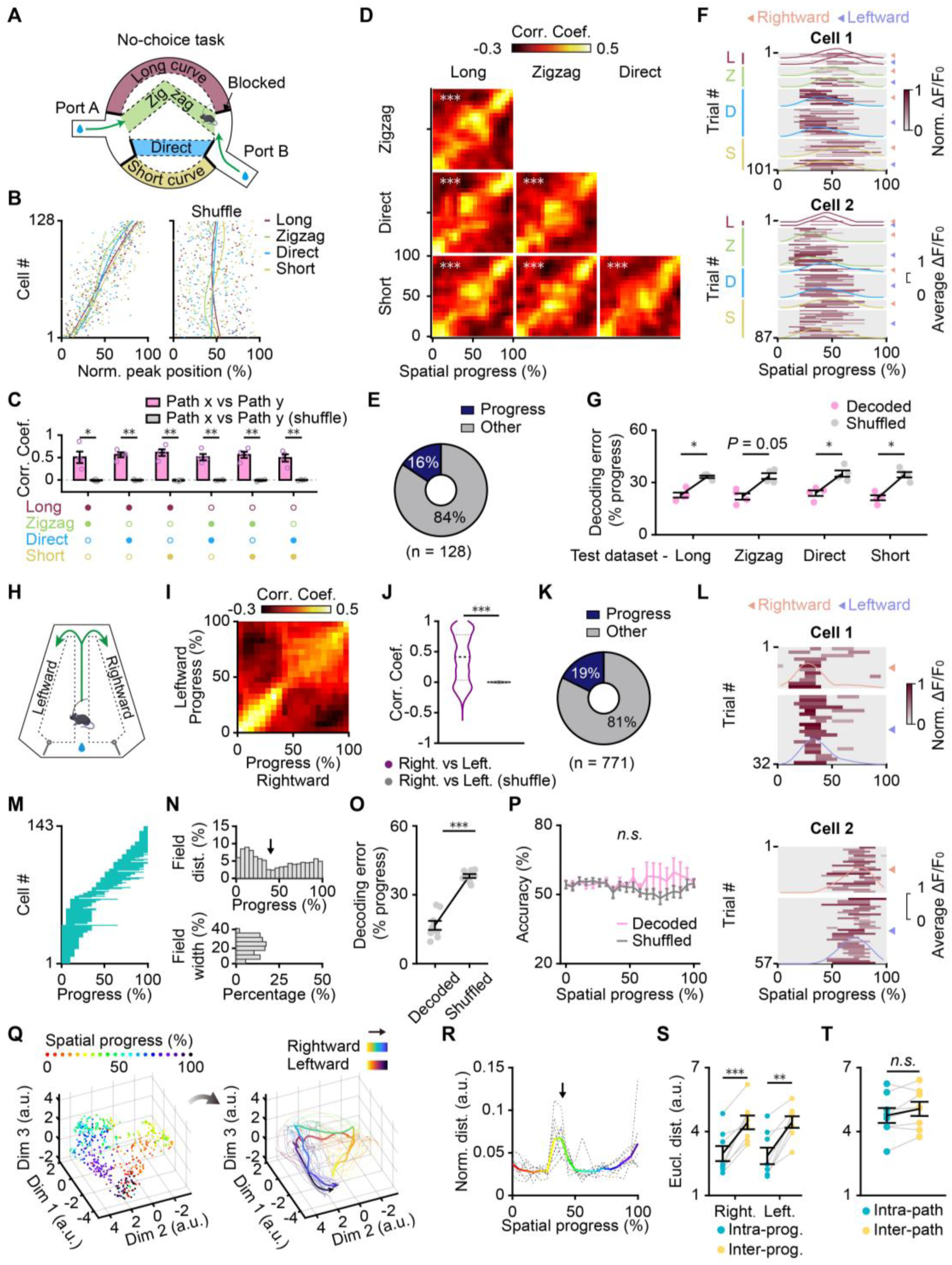
The mPFC encodes spatial progress across different routes and contexts. (A) Schematic showing the modified route choice paradigm with only one path open per session. (B) Left, distribution of normalized peak firing positions of individual neurons sorted by average position across paths. Colored ticks, the average peak firing position of individual neurons in each path. Colored curves, cubic polynomial fits to the peak firing positions in each path. Right, distribution and fitted curve of peak firing positions from the shuffled dataset. (C) Correlation coefficients of normalized peak firing positions between different path pairs (*n* = 4 mice). Filled dots at the bottom indicate the compared path pairs. (D) Population correlation matrices of trial-averaged activity between different path pairs. For each path pair, significance was assessed by comparing the correlation coefficients with those calculated after shuffling neuronal activity in one path (*n* = 128 cells). (E) Proportion of spatial-progress cells in the no-choice task.\ (F) Activity maps of two example spatial-progress cells as a function of spatial progress in the no-choice task. (G) Decoding error for spatial progress calculated using binned neuronal activity from trials in three paths as the training dataset and trials in the other path as the test dataset (*n* = 4 mice). (H) Schematic showing the trapezoid T-maze with geometrically identical paths. Gray lines represent two one-way gates to enforce unidirectional travel. (I) Population correlation matrix of trial-averaged activity between the two paths. (J) Average correlation coefficients calculated from the neuronal activity between the two paths, compared with those from the shuffled dataset (*n* = 771 cells). (K) Proportion of spatial-progress cells in the trapezoid T-maze. (L) Activity maps of two example spatial-progress cells as a function of spatial progress in the trapezoid T-maze. (M, N) Progress fields (M), field distributions, and field widths (N) of spatial-progress cells. The arrow indicates the turn point, which corresponds to 40% progress.(O) Comparison of the decoding error for progress with that calculated from the shuffled dataset (*n* = 8 mice). (P) Decoding accuracy for path types using neuronal activity from individual progress bins (*n* = 8 mice). (Q) Left, neural manifold constructed from both rightward and leftward trials (*n* = 128 cells). Colored dots denote the spatial progress. Right, neural trajectories on the manifold, grouped by paths. Thick arrows, direction of the mean trajectory for each path along spatial progress. Fine lines, neural trajectories of individual trials. (R) Normalized distance between successive bins along the trajectories in (Q). Each dashed gray line represents an animal (*n* = 8 mice). The arrow indicates the turn point, which corresponds to 40% progress. (S) Average Euclidean distance within and between progress stages (*n* = 8 mice). (T) Average Euclidean distance between trajectory pairs of same versus different paths (*n* = 8 mice). \**P*<0.05, \*\**P*<0.01 and \*\*\**P*<0.001. *n.s.*, not significant. Detailed statistics are presented in Table S1. See also Figure S10.

We next asked whether this progress code generalizes across task environments. We imaged mPFC activity in a custom trapezoid T-maze, in which mice chose either a leftward or rightward path to obtain a water reward (Figures 4H and S10F). Progress-aligned activity was highly correlated between leftward and rightward trials (Figures 4I, 4J and S10G). The spatial-progress cells, identified using the same criteria, accounted for approximately 19% of the recorded population and showed nonuniform progress fields with a median width of approximately 23% of the normalized trajectory (Figures 4K–4N and S10H). Notably, progress cells were enriched before the turn point (Figure 4N), possibly reflecting the behavioral salience of this decision point, a pattern not observed in the route choice task (Figure 2K). Consistent with the route choice task, spatial progress could be decoded from mPFC activity above chance, whereas path identity could not be decoded above chance at any progress stage (Figures 4O and 4P). Finally, the population activity was visualized using UMAP, which further revealed structured neural trajectories organized by spatial progress, with similar progress stages clustering together and trajectories from the two paths following comparable ring-like sequential structures (Figures 4Q–4T and S10I). In addition to the larger distances between successive progress bins near trial onset and offset, a distinctive separation was also observed near the turn point (Figure 4R). Together, these results demonstrate that mPFC encodes a generalized representation of spatial progress that is preserved across distinct route configurations and navigational contexts.

### CA1→mPFC inactivation impairs navigational efficiency and progress coding in mPFC

Because spatial-progress recognition during navigation may depend on cognitive map-based self-location, we asked whether hippocampal input to the mPFC contributes to this coding. To selectively manipulate the CA1→mPFC projection while imaging mPFC activity, we expressed hM4Di-mCherry in the ventral hippocampal CA1 neurons projecting to the mPFC and GCaMP6s in mPFC excitatory neurons (Figures 5A and 5B; STAR Methods). After 15 training sessions, mice received three consecutive sessions of saline followed by three consecutive sessions of CNO administration to inhibit the CA1→mPFC projection (Figure 5A). Inactivating the CA1→mPFC projection reduced the number of trials, increased the proportion of random pause, and prolonged route choice time (Figures 5C–5E). In contrast, the path preference, proportion of incomplete trials, and inter-trial interval were not significantly affected (Figures 5F–5I), resembling the behavioral effects of direct mPFC inactivation (Figures 1K–1P). Thus, CA1→mPFC input supports navigational fluency rather than path preference in the route choice task.

**Figure 5.**
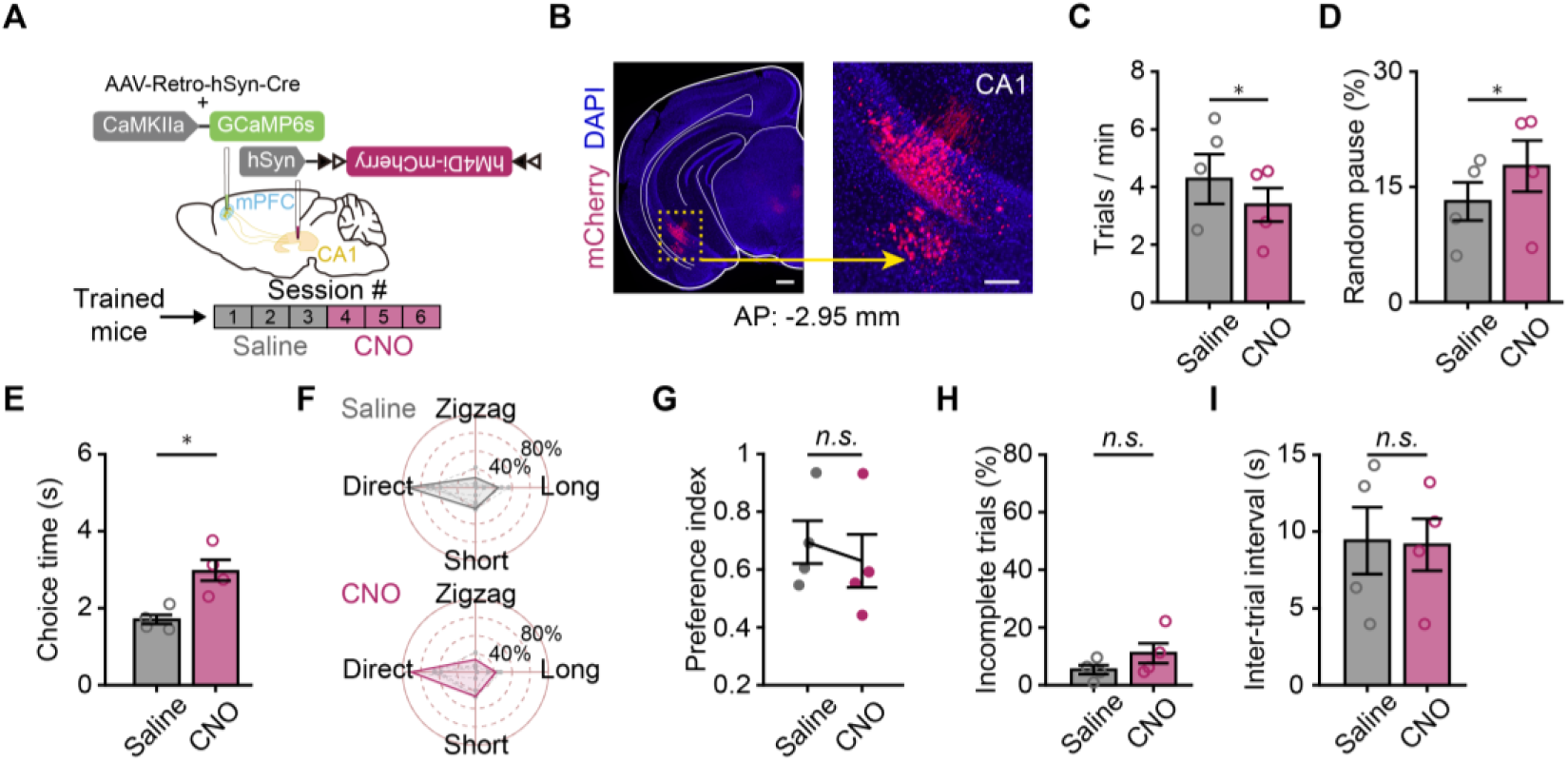
Inactivation of the CA1→mPFC projection impairs the navigational efficiency. (A) Top, schematic of viral injection. Bottom, timeline of the experiment schedule. (B) Representative coronal section of CA1 labeled by hM4Di-mCherry. Scale bars: left: 500 μm, right: 200 μm. (C–E) Comparison of trials per minute (C), proportion of random pause (D), and route choice time (E) before and after inactivation of the CA1→mPFC projection (*n* = 4 mice). (F, G) Comparison of path selection rate (F) and preference index (G) before and after inactivation of the CA1→mPFC projection (*n* = 4 mice). (H, I) Comparison of the proportion of incomplete trials (H) and inter-trial interval (I) before and after inactivation of the CA1→mPFC projection (n = 4 mice). \**P*<0.05. *n.s.*, not significant. Detailed statistics are presented in Table S1. See also Figure S11.

We next assessed whether CA1→mPFC inactivation altered spatial-progress coding in mPFC (Figure 6A). At the population level, progress-aligned activity became less correlated across running directions, across different path types, and across repeated trials of the same path type within and across days (Figures S11A–C). At the single-cell level, some spatial-progress cells maintained similar peak positions after CNO treatment, whereas others showed unstable response profiles across trials and paths (Figure 6B), resulting in higher response jitter and loss of the progress cell identity (Figures 6C and 6D). In addition, the proportion of spatial-progress cells was markedly decreased (Figures 6D and S11D), and approximately half of the previously identified spatial-progress cells lost progress coding following CA1→mPFC inactivation (Figures 6D and 6E). However, the overall distribution and width of progress fields among the remaining neurons were not significantly changed (Figure 6F). These results indicate that CA1→mPFC input is required for stable spatial-progress coding in mPFC.

**Figure 6.**
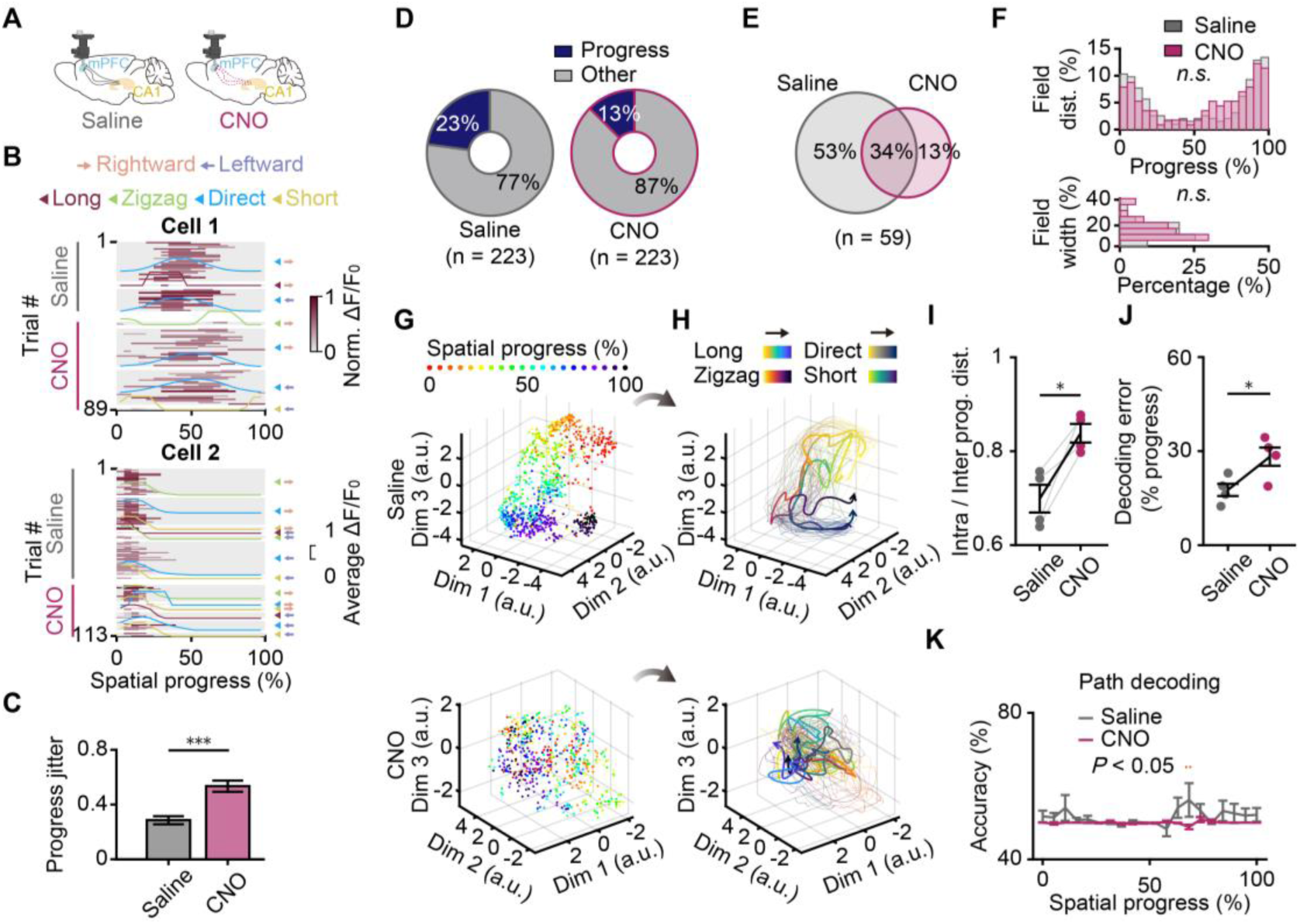
Inactivation of the CA1→mPFC projection impairs the progress coding of mPFC neurons. (A) Schematic of mPFC calcium imaging configuration. Left, saline condition with intact CA1→mPFC projection. Right, CNO condition with chemogenetically inactivated CA1→mPFC projection. (B) Activity maps of two example neurons as a function of spatial progress, which displayed either shifted (top) or stable firing fields (bottom) after inactivation of the CA1→mPFC projection. (C) Comparison of the response jitter for spatial-progress cells before and after inactivation of the CA1→mPFC projection, with activity aligned by spatial progress (*n* = 59 spatial-progress cells identified in either saline or CNO stage). (D) Proportion of spatial-progress cells in mPFC before (left) and after (right) inactivation of the CA1→mPFC projection. (E) Venn diagram illustrating the relationship of spatial-progress cells before and after inactivation of the CA1→mPFC projection.(F) Comparison of the field distributions (top) and field widths (bottom) of spatial-progress cells before and after inactivation of the CA1→mPFC projection (saline, *n* = 63 cells; CNO, *n* = 31 cells). Cells classified as spatial-progress cells for both directions are shown twice. (G) Neural manifold illustrating spatial progress before (top, *n* = 181 cells) and after (bottom, *n* = 144 cells) inactivation of the CA1→mPFC projection. Colored dots denote the spatial progress. (H) Trajectories on the manifolds before (top) and after (bottom) inactivation of the CA1→mPFC projection. (I) Ratio of the Euclidean distances within the same progress stages to distances between them before and after inactivation of the CA1→mPFC projection (*n* = 4 mice). (J) Comparison of the decoding error for spatial progress before and after inactivation of the CA1→mPFC projection (*n* = 4 mice). (K) Comparison of the decoding accuracy for path types before and after inactivation of the CA1→mPFC projection (*n* = 4 mice).\ (G) to (K) are shown for rightward trials (see Figures S11E–I for leftward trials). \**P*<0.05, \*\**P*<0.01 and \*\*\**P*<0.001. *n.s.*, not significant. Detailed statistics are presented in Table S1. See also Figure S11.

Consistently, neural manifold analysis revealed that CA1→mPFC inactivation disrupted the sequential ring-like geometry characteristic of progress trajectories observed during saline sessions (Figures 6G, 6H, S11E, and S11F). Progress stages that were clearly segregated under saline became intermixed after CNO treatment, and the ratio of intra-stage to inter-stage distances increased significantly (Figures 6I and S11G). Decoding error for spatial progress also increased after CNO treatment in both running directions, whereas path decoding remained near chance and was not significantly changed (Figures 6J, 6K, S11H, and S11I). Together, these findings show that CA1→mPFC input is necessary for maintaining a coherent and sequential mPFC representation of spatial progress, thereby supporting fluent goal-directed navigation without directly determining path choice.

## DISCUSSION

Efficient goal-directed navigation requires animals not only to locate themselves in space, but also to monitor their progress from start to goal. However, how the brain constructs and represents such a signal remains unclear. Here, using a custom-designed multi-route navigation paradigm, we identify spatial-progress cells in the mPFC that fire at consistent positions along a normalized start-to-goal axis, whether the path is long or short, straight or curved. Critically, the formation of these cells depends on training and their maintenance relies on the input from the hippocampal CA1. Notably, inactivation of either the mPFC or the CA1→mPFC projection did not alter learned path preference, but decreased the navigational efficiency. These findings suggest that mPFC does not directly compute the optimal route; rather, it represents abstract progress within an ongoing goal-directed episode, thereby supporting fluent navigation across variable routes and contexts.

Spatial-progress cells are distinct from hippocampal place cells, whose firing fields are anchored to absolute spatial positions in the environment(Alme et al., 2014; O’Keefe and Dostrovsky, 1971; Ziv et al., 2013). Although mPFC neurons have also been shown to encode current spatial location(Ma et al., 2023; Zielinski et al., 2019), the spatial-progress cells found in our study cannot be explained by place-specific firing, as they exhibit multiple spatially isolated firing fields that are located at similar spatial-progress stages. More importantly, the number of firing fields equals the number of visited paths. Furthermore, because the lengths and geometries are different for each path and mice dynamically changed the running speed and head direction during traveling at the same progress stage, the activity patterns of these neurons could not be explained by running speed or head direction. An alternative interpretation is that mPFC neurons might encode elapsed time(Heldman et al., 2025) or distance to goal(Howard et al., 2014), as has been described in the hippocampus, rather than spatial progress per se. To exclude this possibility, we aligned the activity of individual neurons with these behavioral variables. When aligned by time of trial onset, mPFC activity shifted progressively toward the trial offset as trial distance increased. Conversely, when aligned by distance to goal, responses shifted to the trial start as trial length increased. In contrast, when aligned by spatial progress, the neurons consistently fired at similar normalized positions across trials of varying durations and lengths. In addition, the response jitter was significantly lower under progress alignment than under the other two alignment conditions. Thus, mPFC neurons represent an abstract progress variable distinct from absolute spatial position and other behavioral correlates of navigation.

Another interesting question concerns the origin of spatial-progress cells in mPFC. From a cognitive perspective, recognizing spatial progress must rely on both task familiarity and perception of external space. For example, estimating progress on a journey from home to the company with multiple possible routes requires not only prior experience with the length and geometry of each route but also knowledge of one’s own position. Unlike hippocampal place cells, which stabilize within minutes after exposure to a novel environment(Priestley et al., 2022), progress-aligned activity of most neurons was disorganized across trials or paths even for several days in the naive stage. The proportion of spatial-progress cells significantly increased only after mice became proficient, along with the formation of sequential neural trajectories in UMAP space. Furthermore, when the CA1→mPFC projection was inactivated, approximately half of the spatial-progress cells lost their coding identity, and the sequential ring-shaped manifold was disrupted. These results support that spatial-progress coding in mPFC is not generated de novo but depends on learning of the task and the transformation of concrete spatial information supplied by the hippocampus.

Our conclusion that inactivation of mPFC impairs navigational fluency and efficiency without altering path preference challenges both our original hypothesis and the well-established role of mPFC in decision making(Euston et al., 2012; Klein-Flügge et al., 2022). Previous studies conducted on the delayed non-match-to-place T-maze task(Benoit et al., 2020; Spellman et al., 2015) have shown that mPFC inactivation leads to decreased choice accuracy in the choice phase. In that task, mice must remember the arm visited in the sample phase and then choose the opposite arm in the choice phase. In contrast, mice in our route choice task develop path preference based on differences in path length and geometry. Moreover, the motivational structure differs between the tasks. The T-maze paradigm imposes more severe penalties for errors (reward omission and time delays), whereas our task entails milder costs for suboptimal choices, primarily in the form of increased energy expenditure and longer travel time associated with longer paths. Last, the post-inactivation difference in choice behavior between the two tasks may arise from distinct memory demands, with our task relying on long-term memory and the T-maze task on short-term working memory. Importantly, our behavioral results align with the evidence that mPFC activity fails to decode the selected path but robustly encodes spatial progress. Together, these results suggest that the mPFC does not affect well-learned, stable path preference in the route choice task.

The mPFC is widely recognized as a key region for higher-order cognitive aspects of spatial navigation. Neurons in this region have been shown to encode basic spatial variables, including current spatial location(Ma et al., 2023; Zielinski et al., 2019) and different trajectory routes(Fujisawa et al., 2008; Ito et al., 2015), akin to hippocampal place cells. Other works have revealed more abstract, task-related representations, including goal locations(Hok et al., 2005; Vogel et al., 2022) and task sequences(Tang et al., 2023). Based on these findings, we propose a conceptual framework in which the mPFC integrates multiple types of navigation-related information, including allocentric spatial location, start and goal locations, structure and proportion of the path traversed, to construct the higher-order cognitive aspects of spatial progress. In addition, previous studies on the HPC-mPFC circuit have demonstrated that the hippocampus primarily encodes the spatial structure of a task, whereas the mPFC learns and implements task rules to guide behavior(Benchenane et al., 2011; Shin and Jadhav, 2016). In particular, the CA1→mPFC projection has been shown to integrate representations of physical and task space, thereby enabling animals to solve challenging spatial alternation tasks(Spellman et al., 2015; Tamura et al., 2017; Van de Maele et al., 2024). Consistent with these findings, our experiments found that inhibiting the CA1→mPFC projection degrades navigational fluency and blurs the spatial-progress coding. These results support that hippocampal-prefrontal communication is essential for maintaining a precise, stable neural code for spatial progress, thereby supporting efficient goal-directed navigation.

Together, our findings identify the mPFC as a core processor of spatial progress, thereby providing a neural mechanism by which animals generate a flexible estimate of “how much have I accomplished” during goal-directed spatial navigation. Moreover, these findings reveal how the brain constructs a flexible readout of ongoing spatial progress, transforming a static representation of the environment into a unified and abstract internal metric of progress. By extracting an invariant measure of progress from diverse experiences (e.g., different directions or paths), this mechanism supports flexible and efficient navigation in complex environments.

## RESOURCE AVAILABILITY

### Lead contact

Requests for further information and resources should be directed to and will be fulfilled by the lead contact, Xinjian Li.

### Materials availability

This study did not generate new unique reagents.

### Data and code availability

The raw datasets for calcium imaging are large (more than 1TB) but are available upon reasonable request. Quantitative data supporting this study are available in Supplementary Information. The custom MATLAB code supporting this study is available in Supplementary Information.

## ACKNOWLEDGMENTS

This work has been supported by Brain Science and Brain-like Intelligence Technology–National Science and Technology Major Project (2022ZD0205000, LXJ; 2021ZD0204101, GLX); the National Natural Science Foundation of China (32371074, LXJ; 32071097, LXJ; 32170991, GLX); National Key Research and Development Program of China (2023YFB4705500, LXJ); MOE Frontier Science Center for Brain Science & Brain-Machine Integration (Zhejiang University, LXJ, GLX); the Fundamental Research Funds for the Central Universities (2023ZFJH01-01 and 2024ZFJH01-01); the Non-profit Central Research Institute Fund of Chinese Academy of Medical Sciences (2023-PT310-01); the Postdoctoral Fellowship Program of CPSF (GZC20241507).

## AUTHOR CONTRIBUTIONS

X.J.L., L.X.G., Y.Z.W., and L.Z.F. conceived the idea and designed the experiments; L.Z.F. and Y.Z.W. performed the surgery; Y.Z.W., W.D.H., L.Z.F., M.X., and S.S.L. performed the experiments and collected the data; Y.Z.W. and M.X. performed the data analysis and drew the figures; X.J.L., L.X.G., Y.Z.W., M.X., L.Z.F., F.F.S., J.T., and B.R.Z. discussed the data and reviewed the manuscript; Y.Z.W. wrote the draft of the manuscript; X.J.L. and Y.Z.W. edited the manuscript; X.J.L. and L.X.G. supervised the project and critically revised the manuscript. All authors approved the final version of the manuscript.

## SUPPLEMENTAL INFORMATION

Document S1. Figures S1–S11. Document S2. Tables S1 and S2. Videos S1 and S2.

## STAR★METHODS

### KEY RESOURCES TABLE

| REAGENT OR RESOURCE | SOURCE | IDENTIFIER |
| --- | --- | --- |
| Bacterial and virus strains |  |  |
| pAAV-mCaMKIIα-GCaMP6s | Shanghai Sunbio Medical Biotechnology | Cat# PMT308 |
| pAAV-hSyn-hM4Di-mCherry | Shanghai Sunbio Medical Biotechnology | Cat# PMT445 |
| AAVretro-hSyn-Cre | Shanghai Sunbio Medical Biotechnology | Cat# PMT387 |
| pAAV-hSyn-DIO-hM4Di-mCherry | Shanghai Sunbio Medical Biotechnology | Cat# PSB1766 |
| Experimental models: Organisms/strains |  |  |
| Mouse: C57BL/6J | Shanghai SLAC Laboratory Animal Co. | RRID: IMSR_JAX:000664 |
| Software and algorithms |  |  |
| MATLAB | Mathworks | https://www.mathworks.com/products/matlab.html |
| ImageJ | National Institutes of Health | https://imagej.nih.gov/ij/index.html; RRID: SCR_003070 |
| NoRMCorre toolbox | Pnevmatikakis and Giovannucci, 2017 | https://github.com/flatironinstitute/NoRMCorre |
| CellReg toolbox | Sheintuch et al., 2017 | https://github.com/zivlab/CellReg |
| CellSort toolbox | Mukamel et al., 2009 | https://www2.mathworks.cn/matlabcentral/fileexchange/25405-emukamel-cellsort |
| Custom MATLAB scripts for data analysis | This work | N/A |
| Other |  |  |
| Stereotaxic apparatus | RWD Life Science | https://www.rwds.com/product-solutions/life-sciences/modeling/stereotaxic/data_20.html |
| Syringe pump | KD Scientific | https://www.kdscientific.com/legato-130-syringe-pump.html |
| Miniature microscope | Inscopix | https://inscopix.com/nvoke-system/ |
| GRIN lens | Inscopix | Cat# 1050-004605 |

### EXPERIMENTAL MODEL AND SUBJECT DETAILS

A total of 31 adult male C57BL/6J mice (8–10 weeks, 20–25 g) were used in this study. Mice were housed 4–5 per cage at a room temperature of 25 ± 1 °C and a humidity of 40 ± 5% on a 12-h light/dark cycle (lights on at 07:00 and off at 19:00). After surgery and during behavioral training, mice were housed individually. Mice were provided food and water ad libitum until the behavioral test. The behavioral test and calcium imaging were conducted during the daytime. All mice performed the route choice task. Among the 31 mice, 16 mice were used exclusively for behavioral test after chemogenetic inactivation of mPFC (8 received saline, 8 received CNO); 11 mice were used for both behavioral test and calcium imaging; 4 mice were used to assess behavioral and neural activity changes before and after inactivation of the CA1→mPFC projection. Among 15 calcium imaging mice, 12 mice performed the trapezoid T-maze. All experiments were conducted in accordance with the guidelines for the care and use of laboratory animals at Zhejiang University and were reviewed and approved by the Animal Advisory Committee at Zhejiang University (ZJU20260046).

### METHOD DETAILS

#### Virus injection

The viral injection protocol was based on previous methods with some modifications(Lyu et al., 2022; Zhao et al., 2022). The surgical tools and stereotaxic apparatus were sterilized with 70% ethanol 30 min before surgery. Mice were anesthetized with Avertin (1.5% solution, 0.01 ml/g, i.p.) and pretreated with dexamethasone (0.2 mg/kg, s.c.) before surgery to mitigate inflammation. The fur between the eyes and ears was trimmed, and the skin was disinfected with 70% ethanol using three swabs. The eyes of the mice were protected with ophthalmic lubricant. To alleviate acute pain during surgery, 2% lidocaine was administered locally (s.c.). Mice were placed in a stereotaxic apparatus (RWD Life Science) with the skull leveled along both the anteroposterior and mediolateral axes. A small craniotomy (∼1 mm diameter) was made with a high-speed micro-drill over the target region. Viral vectors were delivered in a volume of 600 nl per injection at a rate of 30 nl/min using a syringe pump (KD Scientific). For mice used for calcium imaging, pAAV-mCaMKIIα-GCaMP6s (Shanghai Sunbio Medical Biotechnology) was injected unilaterally into the right mPFC (AP: +1.94 mm, ML: +0.50 mm, DV: +2.10 mm). All AP and ML coordinates are relative to bregma, and DV coordinates are relative to the brain surface.

### Behavioral training

Before behavioral training, mice were handled for at least 5 days (5 min per day). Mice used for calcium imaging underwent another 5 days of habituation to the head-mounted microscope (5 min per day). Baseline body weight was then recorded, and mice were water-restricted 1 day before training. Water restriction was temporarily suspended if body weight fell below 80% of the baseline. A 15-day training and testing protocol was used in our experiments (15 min/session, 1 session/day). Behavior was recorded with a top-view camera (1080p, Ordro) at a frame rate of 30 Hz. All behavioral experiments were conducted during the daytime (from 2 p.m. to 5 p.m.). After training, the arena was cleaned thoroughly with 70% ethanol, and the water ports were cleaned and visually inspected. The movement of mice was tracked using custom MATLAB code by contrasting the black fur against the white maze. Aberrant movement displacements in some frames were manually corrected. The movement trace of each animal was plotted and verified by a second experimenter.

### Route choice paradigm

Spatial navigation in a natural environment often involves multiple routes to reach the destination. To better understand the neural mechanisms underlying route choice behavior, a custom route choice paradigm was designed (Figure 1A). The maze was constructed from white acrylic (3.2 mm thick), which comprises a central circular arena (60 cm radius) and two rectangular boxes (25 cm × 10 cm). Each box is equipped with a water-delivery port (port A or B) located on the wall opposite the arena. In addition, four distinct paths with different geometries are arranged on the central circular arena, connecting the two reward boxes. These paths are named as follows: a long curve, a zigzag path, a direct path, and a short curve, with canonical trajectory lengths of 210 cm, 165 cm, 135 cm, and 153 cm, respectively. The four paths vary in length and geometry (e.g., the direct path was the shortest and straightest, whereas the long curve was the longest and most circuitous). The width of both boxes and paths is 10 cm, and the height of the walls is 15 cm. Water delivery was controlled by a microcontroller and triggered by lick detection via a infrared sensor. Mice received a water reward only upon completing a full traversal to the opposite port; returning to the same port without reaching the opposite side did not trigger reward delivery.

Mice were motivated to select different paths and navigate back and forth between two water ports. A trial was defined as a complete traversal from one port to the other. Only trials performed without backtracking or retreating were classified as complete trials and used for analysis. For simplicity, trials from port A to port B and those from port B to port A were referred to as rightward and leftward trials, respectively. Trials, running direction, and chosen path were automatically identified using custom code based on movement tracking.

For each trial, six behavioral events were extracted using custom code: (1) trial onset (initiation of movement after reward consumption), (2) leave reward (exit from the start box), (3) path start (entry into the chosen path), (4) path end (exit from the path), (5) enter reward (entry into the reward box), and (6) trial offset (arrival at the reward port). Event timestamps were manually confirmed by a second experimenter.

Behavioral performance was quantified using several behavioral parameters: proportion of incomplete trials, route choice time, proportion of random pause, path selection rate, and path preference index. The proportion of incomplete trials was calculated using the following formula:

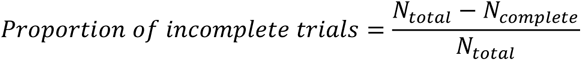

where *N_complete_* denotes the number of complete trials, and *N_total_* represents the total number of trials. Route choice time was defined as the average time mice spent in the route selection regions. The proportion of random pause was calculated from the speed profile of each mouse. To do so, the instantaneous speed was derived from the movement trace and smoothed with a 0.5-s moving-average filter. Periods within a trial when instantaneous speed fell below 1 cm/s were classified as random pauses. The proportion of random pause time was calculated as the ratio of random pause duration to trial duration. The selection rate for each path was computed as the fraction of complete trials on that path relative to all complete trials. Finally, a preference index was adapted to qualify path preference:

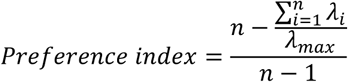

where *n* denotes the number of available paths (*n* = 4), *λ_i_* denotes the number of complete trials on path *i*, and *λ_max_* denotes the maximal number of complete trials across all paths. The preference index ranges from 0 (all paths chosen equally) to 1 (only a single path chosen in a session).

#### Trapezoid T-maze paradigm

A T-maze with a trapezoidal layout was constructed from white acrylic (3.2 mm thick) following previous studies with some modifications(Xiang et al., 2019). The maze consists of a main stem (80 cm), two choice arms (each 90 cm, angled 20° relative to the stem), and two return arms (20 cm) (Figure 4H). A water-delivery port is located at the start point of the main stem. One-way gates are mounted on each return arm near the reward port to enforce unidirectional navigation. After consuming water, the mouse can only run forward along the main stem, choose one of the two choice arms, and return via the connected return arm. Revisiting the reward port without completing a full loop does not trigger water delivery. Lick detection and water delivery were implemented similarly to those in the route choice paradigm. Trials were automatically identified using custom code. Behavioral performance was assessed with parameters analogous to those used in the route choice paradigm.

#### Calcium imaging in freely moving animals

##### GRIN lens implantation

The procedures for gradient refractive index (GRIN) lens (Inscopix, 1.0 mm diameter, 4.0 mm length) implantation and baseplate attachment were similar to those described in previous works(Kingsbury et al., 2019; Liang et al., 2018; Murano et al., 2022). Two weeks after virus injection, a 1-mm diameter GRIN lens was implanted into the mPFC under anesthesia. In brief, a small craniotomy (∼1.1 mm diameter) was first made over the target coordinate (AP: +1.94 mm, ML: +0.50 mm). Then, to reduce pressure on the brain tissue during lens insertion, an insertion tract (∼1.8 mm ventral to the brain surface) was created using a 19-gauge blunted needle (∼1 mm diameter), and the brain tissue above the mPFC was slowly aspirated along the tract. The GRIN lens was then slowly lowered into the cavity until its tip reached the target depth (∼1.9 mm ventral to the brain surface) and secured to the skull using dental cement. After implantation, the top of the lens was covered with elastomeric impression material for protection. Three weeks after lens implantation, the mice were anesthetized again, and a baseplate was installed above the GRIN lens to attach the microscope for calcium imaging. In detail, the impression material was first removed. Then, the baseplate was attached to the miniature microscope (Inscopix, nVoke 2.0) and lowered to the top of the lens. When clear cell bodies of imaging neurons were visible in the field of view, the baseplate was fixed to the skull using dental cement. Before each imaging session, the microscope was reattached and secured to the baseplate via a side-locking screw. This approach enabled us to concurrently record the calcium activity from hundreds of excitatory neurons in the mPFC of freely moving mice.

##### Calcium imaging

Calcium imaging was performed during the behavioral task via a head-attached microscope (LED power: 0.1–0.6 mW; camera resolution: 1280 × 800 pixels)(Lyu et al., 2022). Images were acquired at a frame rate of 30 Hz via Inscopix data acquisition software (field of view: 1050 μm × 650 μm). Before each imaging session, the lens and baseplate were carefully cleaned with lens paper. The microscope was then attached to the baseplate and the focal plane was adjusted until clear somatic signals were observed. The behavioral video was synchronized with the calcium imaging stream via the trigger-out signal from nVoke HD.

##### Calcium event detection

Calcium activity was extracted using custom scripts in MATLAB following published work(Lyu et al., 2022; Mukamel et al., 2009; Wang et al., 2017). Raw videos were first spatially down-sampled fourfold along spatial dimensions and then motion-corrected with the stabilizer plugin in ImageJ (National Institutes of Health) or the NoRMcorre toolbox(Pnevmatikakis and Giovannucci, 2017) to compensate for translational drift. The mean fluorescence intensity of each pixel in each session (15 min) was calculated as the baseline (F_0_), and fluorescence changes in pixel intensity at time t were expressed as (F_t_ – F_0_)/F_0_ or ΔF/F_0_. Active neuronal calcium signals were derived via principal component and independent component analysis (PCA-ICA) implemented with the CellSort and fastICA toolboxes. This method decomposes the spatiotemporal ΔF/F_0_ matrix into independent components based on the skewness of the data distribution. Each component comprises a characteristic spatial filter over the imaged area and a corresponding temporal signal during the imaging period. The spatial filter and the temporal signal of each component were graphed and inspected by investigators who were blind to the experimental conditions of each video. If the spatial filter for a component overlapped with the dark shadows cast by blood vessels in the F_0_ image, this component was likely contributed by blood flow and was therefore rejected. Furthermore, because calcium signals exhibit a characteristic fast-rising and slow-decaying time course, the temporal skewness of calcium signals was expected to be positive, and those components with skewness <1 were rejected. For each selected component, neuron centroids were defined as the brightest 3 × 3-pixel region within each spatial filter. The corresponding temporal signal of each selected component was calculated from the ΔF/F_0_ video by subtracting the median value of the background area (outside the cell body) from the average value of the cell area (Figures S2D and S2E).

To identify periods of elevated neuronal activity, the rising phase of each calcium event (peak ΔF/F_0_ > 3 standard deviations of baseline fluctuation) was searched for. The onset of this rising phase was defined as the time point when the first derivative of the ΔF/F_0_ trace (calculated in a 200-ms moving window) rose above 0 and continued to increase above 5 standard deviations of baseline fluctuation; and the offset of this rising phase was defined as the time point where the first derivative fell below 0.

##### Identify the same neurons across sessions

To track the same neurons across imaging sessions, we applied the open-source cross-session registration algorithm CellReg(Sheintuch et al., 2017). This method compares the spatial correlations of nearby cells to determine whether highly correlated footprints proximal in the field of view are probabilistically assigned as the same cell. All putative matched neurons generated by CellReg were manually verified. If clear misalignment (e.g., non-overlapping somatic regions, inconsistent morphology, or abnormally low cross-session matching rate) was found during visual inspection, we iteratively adjusted the registration parameters—including ‘alignment_type’, ‘reference_session_index’, and ‘p_same_threshold’—until the spatial correspondence across sessions was optimized. This combined approach ensured that only consistently identified neurons were used for subsequent analyses.

##### Chemogenetic manipulation

To directly inactivate the mPFC, pAAV-hSyn-hM4Di-mCherry (Shanghai Sunbio Medical Biotechnology) was injected bilaterally into the mPFC (AP: +1.94 mm, ML: ±0.50 mm, DV: +2.10 mm). Mice were randomly assigned to experimental (CNO) or control (saline) groups. Saline or CNO (3 mg/kg, i.p.; Sigma-Aldrich) was administered before each session throughout training. To inactivate the CA1→mPFC projection, multi-site injections were performed following a previously established protocol(Spellman et al., 2015). First, AAVretro-hSyn-Cre and pAAV-mCaMKIIα-GCaMP6s were co-injected into the right mPFC (AP: +1.94 mm, ML: +0.50 mm, DV: +2.10 mm). Then, pAAV-hSyn-DIO-hM4Di-mCherry was injected bilaterally into the CA1 area of the hippocampus (4 sites; AP: −2.95 mm, ML: ±3.15 mm, DV: +4.30 mm; and AP: −3.15 mm, ML: ±3.00 mm, DV: +3.90 mm). The viral injection volume was 600 nl per site, and the injection rate was 30 nl/min. After 15 training sessions, mice received three consecutive sessions of saline administration, followed by three consecutive sessions of CNO administration (3 mg/kg, i.p.).

##### Trajectories linearization and spatial segmentation

To better quantify progress-related neuronal activity, we developed a pipeline to transform the curved movement trace into linearized positions along spatial progress. To do so, we derived a standard trial trajectory for each path, and the movement traces of individual trials were projected onto their corresponding standard trajectories to obtain normalized positions along spatial progress. In detail, the movement traces of mice were first tracked. Second, the movement traces were resampled using piecewise cubic Hermite interpolating polynomial (PCHIP) interpolation and normalized from 0 to 1 using the cumulative path length. Third, the standard trial trajectory for each path was derived by averaging the normalized movement traces in each path and smoothed using a smoothing spline interpolation (λ = 0.9). Last, the movement trajectory in each trial was projected onto its corresponding standard trajectory by computing the shortest Euclidean distance to all points on the standard trajectory. With this method, we mapped the mice’s two-dimensional movement in different paths onto a one-dimensional axis based on the relative spatial progress (Figure S2H).

To compare the neural activity across geometrically distinct paths, the standard trajectory of each trial was segmented into 20 spatial bins: 2 bins covered the start box, 16 bins spanned the path, and 2 bins covered the end box (Figure S2G). With this method, we reconceptualized this linearized axis in terms of spatial progress at a 5% step, where 0% corresponds to trial onset and 100% to trial offset. This allowed us to express the neural activity as a function of spatial progress, independent of the geometry or length of the path.

#### Quantifying the correlation of neuronal activity between directions and between paths

##### Between directions

To quantify the activity correlation between two running directions, we constructed trial-averaged activity profiles by averaging the progress-aligned calcium activity across all trials in each running direction. Pearson correlation coefficients were then calculated between rightward and leftward trials, between rightward and spatially reversed leftward trials, and between rightward and shuffled leftward trials (1,000 times random permutations).

##### Between paths

To exclude the influence of running direction, correlation coefficients between path types were computed separately for rightward and leftward trials. The trial-averaged activity profiles for each path were constructed by averaging the progress-aligned calcium activity across all trials completed on that path. Pearson correlation coefficients were then calculated for each pair of paths.

##### Mapping the activity profiles of single neurons under different alignment conditions

To characterize the activity pattern of individual neurons, we aligned the activity of individual trials by time from trial onset, distance to goal, and normalized distance from onset to offset (i.e., spatial progress). Trials were separated by path type and running direction. Average response curves for each path type and running direction were overlaid on the activity maps.

##### Quantifying the response jitter across different alignment conditions

To assess the response consistency, we computed the response jitter of individual neurons under three alignment conditions. In detail, we first identified the neuronal peak firing positions of individual trials at a given alignment condition (time from trial onset, distance to goal, or normalized distance from onset to offset). We then calculated the standard deviation of peak positions across trials. To correct for scale differences across different alignment conditions, we calculated the chance level of standard deviation using the peak firing positions from the shuffled dataset. Finally, we defined the response jitter for each neuron as the ratio of the observed standard deviation to the chance level. Lower jitter indicates higher trial-to-trial consistency in the timing or location of peak firing.

##### Identification of spatial-progress cells

To identify the spatial-progress cells, we applied the following selection criteria (Figure S5A): (1) exhibited place-cell-like spatial tuning within individual paths(Fischler-Ruiz et al., 2021). For each path, a neuron had at least one bin with trial-averaged activity more than 3 times the trial-averaged activity over the entire trajectory, and z-scored activity greater than 1 in more than 25% of trials; (2) fired at similar spatial-progress stages across geometrically distinct paths. The difference in the peak firing positions between different paths was less than 25% of the full progress scale.

For each qualified neuron, we further calculated the progress field following previous studies of place field with some modifications(Ziv et al., 2013). To do so, we first calculated the average response curve as a function of spatial progress for each path. The overlapping region of these curves was then identified. Within this overlapping region, the progress field was defined as the range of bins where the response exceeded half of the maximum value.

##### Low-dimensional manifold visualization with UMAP

To characterize how spatial progress is represented at the population level in the mPFC, we performed unsupervised dimensionality reduction on the population activity vectors using Uniform Manifold Approximation and Projection (UMAP) following previous studies(Tang et al., 2023; Zutshi et al., 2025). For each trial, a population vector was constructed by averaging the calcium activity of all simultaneously recorded neurons within each progress bin. UMAP was applied separately to trials from each running direction or from all complete trials. Hyperparameters were set as ‘n_components’ = 3, ‘n_neighbors’ = 50, ‘min_dist’ = 0.6, ‘metric’ = ‘cosine’, and ‘init’ = ‘spectral’. In the resulting low-dimensional manifold, each dot corresponds to the population activity of a single spatial-progress bin from a single trial, colored by progress (0%–100%). The performance was quantified by comparing the Euclidean distances between dots of the same progress bins (intra-progress stage distance) to those of different progress bins (inter-progress stage distance). Only animals with at least 50 reliably registered neurons were included in UMAP and decoding analyses.

To further examine the evolution of population activity along spatial progress within individual trials, we connected dots from the same trials in order of spatial progress to form single-trial neural trajectories. Neural trajectories were then averaged within each trial group (chosen path or running direction) and were color-coded by both group identity and progress stage. Last, the neural trajectories were smoothed by cubic spline interpolation. The Euclidean distances between successive progress bins along each trajectory were calculated and normalized by the total trajectory length. We then quantified the geometric organization of these manifolds by comparing the average pointwise Euclidean distances between single-trial trajectories of the same path or direction to those of different paths or directions.

##### Decoding the spatial progress using a support vector machine

To decode the spatial progress, we employed a linear support vector machine (SVM) algorithm using binned neuronal activity following previous works(Curreli et al., 2022; Tsutsui et al., 2016). Decoding was performed only on the dataset from the trained stage, as the number of trials in the naive stage was insufficient for robust model training. In addition, to recruit enough trials, we pooled trials from three consecutive sessions together for subsequent decoding. We decoded the spatial progress separately for each running direction. In detail, trials for each running direction were randomly partitioned into training (75%) and test (25%) datasets. A linear support-vector regression model (MATLAB fitrsvm, linear kernel) was then trained to decode the spatial progress using population activity vectors from the training dataset, and the decoder was used to decode the spatial progress using the test dataset. The decoding error was calculated as the root mean square error (RMSE) between the predicted and actual spatial progress (0–100%, trial onset to offset). 10-fold cross-validation was used to reduce the decoding bias. The chance level was calculated by shuffling the population vectors 1,000 times.

We also trained the decoder using the dataset from the most frequented path and evaluated the decoding accuracy on the second most frequented path. Additionally, we evaluated the stability of progress decoding by training the decoder on the first half of the trials and testing the decoding on the second half, or vice versa. To exclude the possibility that decoding accuracy was driven primarily by neural activity at trial boundaries, we further tested the decoding accuracy using only the progress from 10% to 90%.

##### Decoding the running direction or path using a naive Bayes classifier

To decode the running direction or chosen path, we used a naive Bayes classifier based on kernel density estimation. All decoding was carried out exclusively on data from the trained stage.

Because the number of trials across different paths was imbalanced after training, random oversampling was applied to equalize the trial numbers following previous studies(Chawla, 2005). In addition, if the number of trials for the long curve was less than 3, the long curve was excluded from decoding for that animal. We then implemented a naive Bayes classifier (MATLAB ‘fitcnb’, ‘DistributionNames’ set to ‘kernel’) to decode path type or running direction for each trial at each progress bin using the population calcium activity of all recorded neurons (75% of trials were used for training and 25% of trials for testing). 10-fold cross-validation was used to reduce the decoding bias. Chance level was calculated from shuffled dataset (randomly permuting the neuron sequence 1,000 times). Statistical significance was determined by comparing the decoding accuracy to the chance level.

##### Histology

The expression of GCaMP6s and hM4Di-mCherry in mPFC, as well as retrograde labeling of hM4Di-mCherry in hippocampal CA1, was histologically confirmed in fixed brain tissue from the mice after experiments(Lyu et al., 2022; Solstad et al., 2008). Mice were deeply anesthetized (sodium pentobarbital; overdose) and then transcardially perfused with saline followed by 4% paraformaldehyde [PFA; in 0.01 M phosphate-buffered saline (PBS), pH 7.4]. Brains were post-fixed in 4% PFA (24 h at 4°C) and then transferred to 15% and 30% sucrose solutions (24 h each at 4°C). Serial coronal sections (30 µm) were prepared using a cryostat microtome (Leica CM1950, Germany). Brain sections were washed with PBS and mounted on the coverslips with DAPI-containing fluorescent mounting medium (Solarbio). The injection sites and labeling efficiency were further confirmed via fluorescence imaging performed on a Leica SP8 confocal laser-scanning microscope.

### QUANTIFICATION AND STATISTICAL ANALYSIS

Detailed statistical information for all experiments can be found in Tables S1 and S2. All statistical analyses were performed using MATLAB R2023a (Mathworks). Data were presented as mean ± s.e.m. unless otherwise specified. The normality assumption was tested with the Shapiro-Wilk test. In all figures, * represented *P* < 0.05, ** represented *P* < 0.01, *** represented *P* < 0.001, and *n.s.* represented not significant (*P* > 0.05). Parametric tests (independent t-test, paired t-test, one-way ANOVA, or one-way repeated-measures ANOVA) were used when data met normality assumptions; otherwise, non-parametric equivalents were applied (Mann–Whitney U test, Wilcoxon signed rank test, or Kolmogorov-Smirnov test). Post hoc comparisons for multiple-group data were performed using the Bonferroni correction. All tests were two-sided. Pearson’s correlation was used for calculating the correlation coefficients, and random permutations were used for determining the chance level.

