## Supplementary Figures for "A Generalized Spatial Progress Code for Navigation in the Medial Prefrontal Cortex"

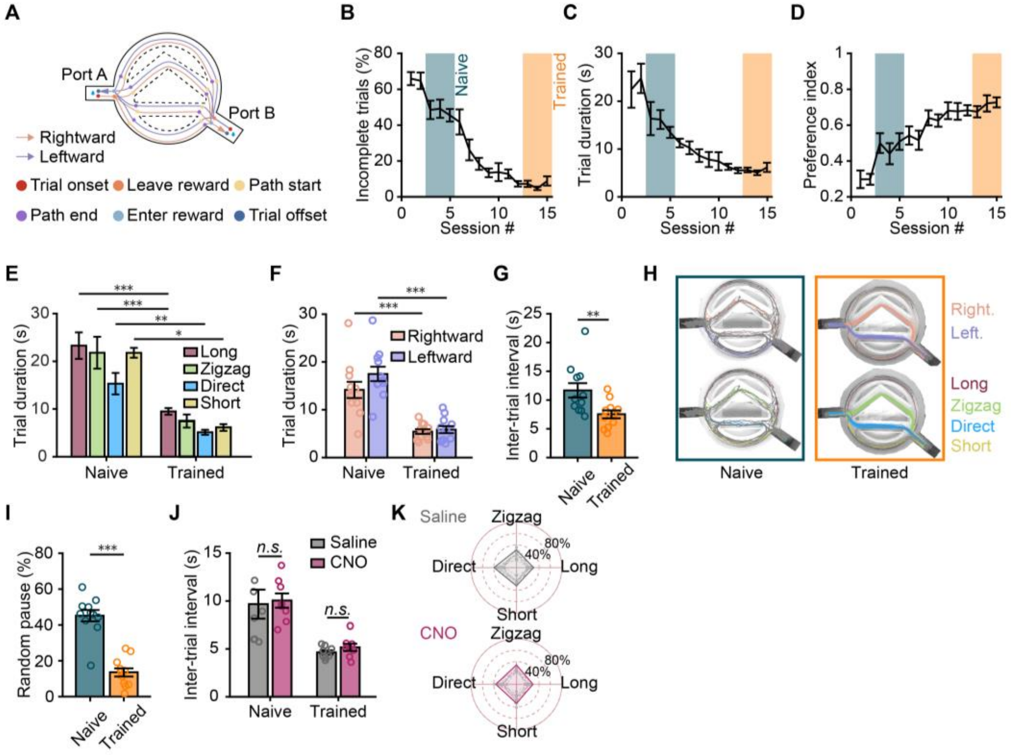


**Figure S1. Training improves the navigational performance in the route choice maze. Related to Figure 1.**

(A) Pseudo-movement trajectories in the route choice maze showing rightward (light pink) and leftward (light purple) trials. The positions of six behavioral events are annotated along the trajectories: trial onset, leave reward, path start, path end, enter reward, and trial offset.

(B–D) Proportion of incomplete trials (B), trial duration (C), and path preference index (D) across 15 training sessions (*n* = 11 mice).

(E–G) Comparison of trial duration for individual paths (E), trial duration for individual directions (F), and inter-trial interval (G) between the naive and trained stages (*n* = 11 mice).

(H) Movement trajectories of an example mouse in the naive and trained stages, color-coded by running direction (top) or by chosen path (bottom).

(I) Comparison of the proportion of random pause between the naive and trained stages (*n* = 11 mice).

(J) Comparison of inter-trial interval between the saline and CNO groups (*n* = 8 mice per group).

(K) Path selection rates of the saline and CNO groups in the naive stage (*n* = 8 mice per group).

*$P$<0.05, **$P$<0.01 and ***$P$<0.001. *n.s.*, not significant.

Detailed statistics are presented in Table S2.

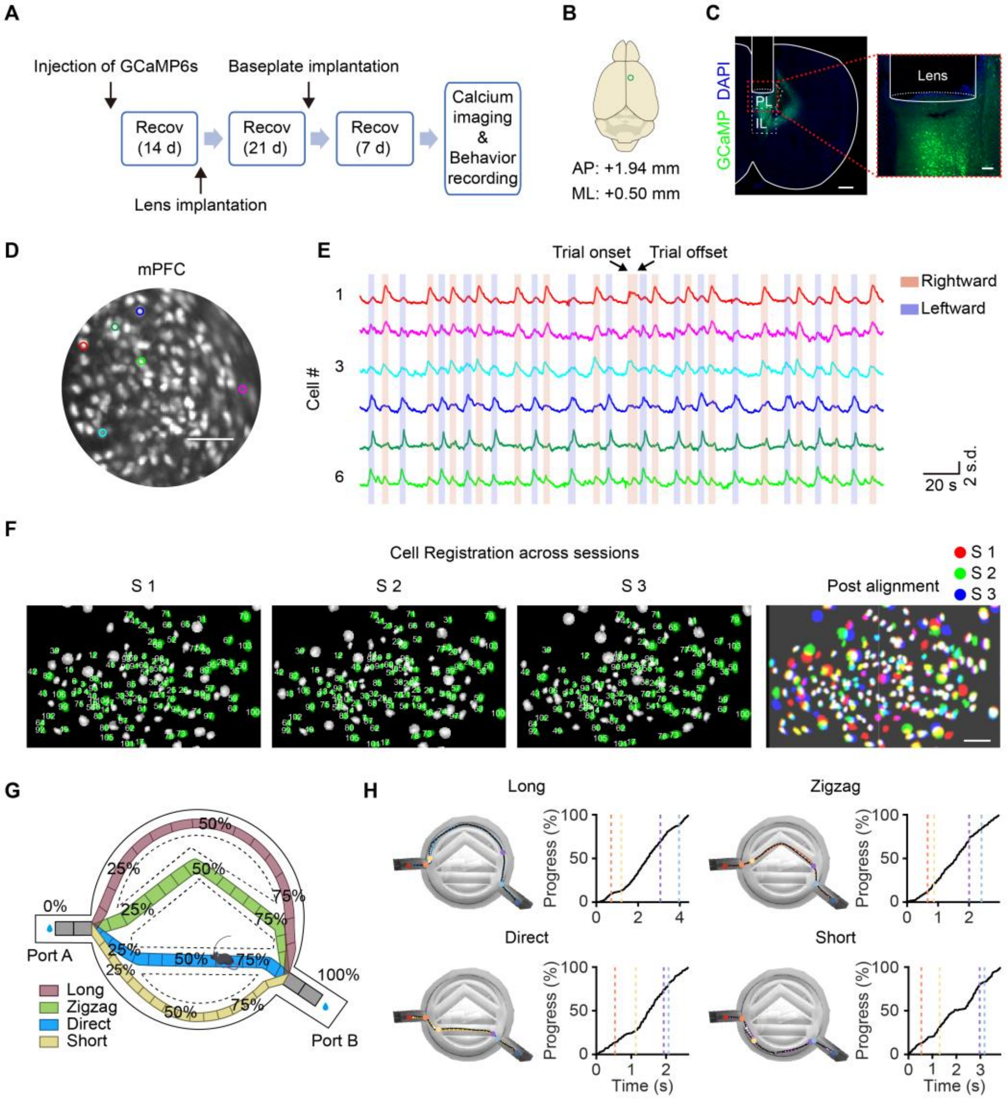


**Figure S2. Imaging and analysis of mPFC calcium activity during navigation in the route choice maze. Related to Figure 2.**

(A) Outline of the experimental procedure.
(B) Virus injection site.
(C) Left, coronal brain section showing the position of GCaMP6s expression and GRIN lens implantation. Scale bar, 500 μm. Right, enlarged view of the red box in the left panel. Scale bar, 100 μm. PL, prelimbic cortex; IL, infralimbic cortex.
(D) Representative maximum projection fluorescence image of mPFC excitatory neurons across 27,000 $\Delta F/F_0$ frames. Scale bar, 100 μm.
(E) Example traces showing the $Ca^{2+}$ transients of individual neurons from regions of interest in (D) (color-matched).
(F) Left, representative images showing detected cells from three consecutive sessions (S1–S3) in an example mouse. Colored circles denote the same cells registered across sessions. Right, overlay of identical cells registered across sessions (white) using the CellReg package. Scale bar, 100 μm.

(G) Schematic showing the spatial-progress bins along the linearized trajectories for four different paths. For each path, the movement trajectory of each trial was projected onto a standard trajectory that was linearly segmented into 20 bins (2 bins for the start box, 16 bins for the central path, and 2 bins for the end box) to obtain spatial progress. The scale was stepped by 5% progress.

(H) Example movement trajectories from each of the four paths and the corresponding spatial progress as a function of elapsed time. Colored dots and dashed lines indicate six behavioral events: trial onset (red), leave reward (orange), path start (yellow), path end (purple), enter reward (light blue), and trial offset (dark blue).

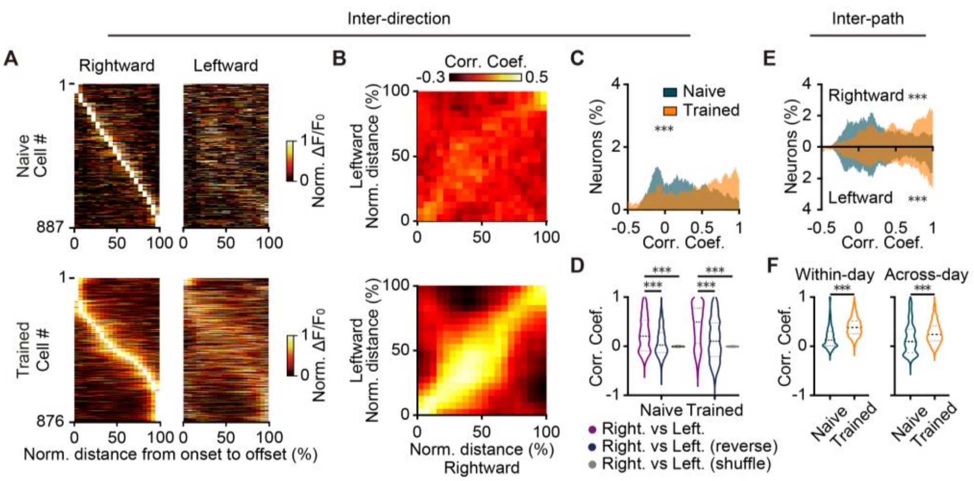


**Figure S3. Training enhances the correlation of mPFC activity across running directions and paths. Related to Figure 2.**

(A) Heatmaps showing trial-averaged neuronal activity aligned by normalized distance from trial onset to offset in the naive (top) and trained (bottom) stages. Left, rightward trials; right, leftward trials. Neurons for both rightward and leftward trials are sorted by peak firing position along spatial progress in rightward trials. Only neurons active in both directions are shown.

(B) Correlation matrices between trial-averaged activity of rightward and leftward trials. Top, the naive stage; bottom, the trained stage.

(C) Distributions of correlation coefficients between trial-averaged activity of rightward and leftward trials (naive, $n$ = 1023 cells; trained, $n$ = 891 cells).

(D) Comparison of correlation coefficients between rightward and leftward trials with those after reversing or shuffling neuronal activity of leftward trials (naive, $n$ = 1023 cells; trained, $n$ = 891 cells).

(E) Distributions of correlation coefficients between trial-averaged activity of different paths, computed separately for rightward (top) and leftward (bottom) trials to exclude the influence of running direction (naive, $n$ = 1023 cells; trained, $n$ = 891 cells).

(F) Comparison of correlation coefficients for the same path type between the naive and trained stages (naive, $n$ = 1023 cells; trained, $n$ = 891 cells). Left, within days; right, across days.

***$P$<0.001.

Detailed statistics are presented in Table S2.

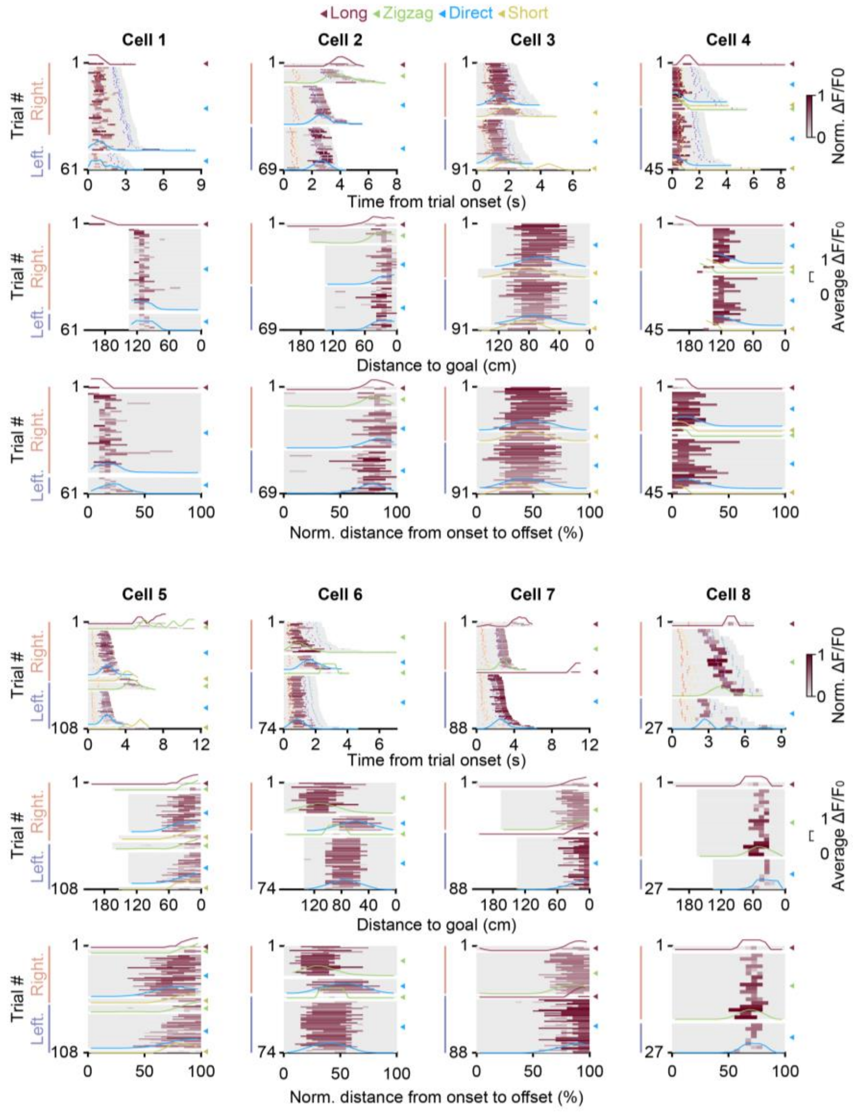


**Figure S4. Comparison of neuronal activity across different alignment conditions. Related to Figure 2.**

Activity maps of example mPFC neurons as a function of time from trial onset (top), distance to goal (middle), or normalized distance from onset to offset (bottom). Overlaid curves show the average activity calculated separately from trials of each path type. Colored dashed lines mark behavioral events: leave reward (orange), path start (yellow), path end (purple), and enter reward (light blue). The response profiles differed depending on the alignment conditions. When aligned by time from trial onset, responses shifted toward trial offset as trial duration increased. When aligned by distance to goal, responses shifted toward trial onset as path distance increased. In contrast, when aligned by spatial progress (normalized distance from onset to offset), each neuron discharged at similar positions across trials, irrespective of path geometry or movement direction. Individual neurons exhibited distinct firing peaks distributed at different positions along spatial progress.

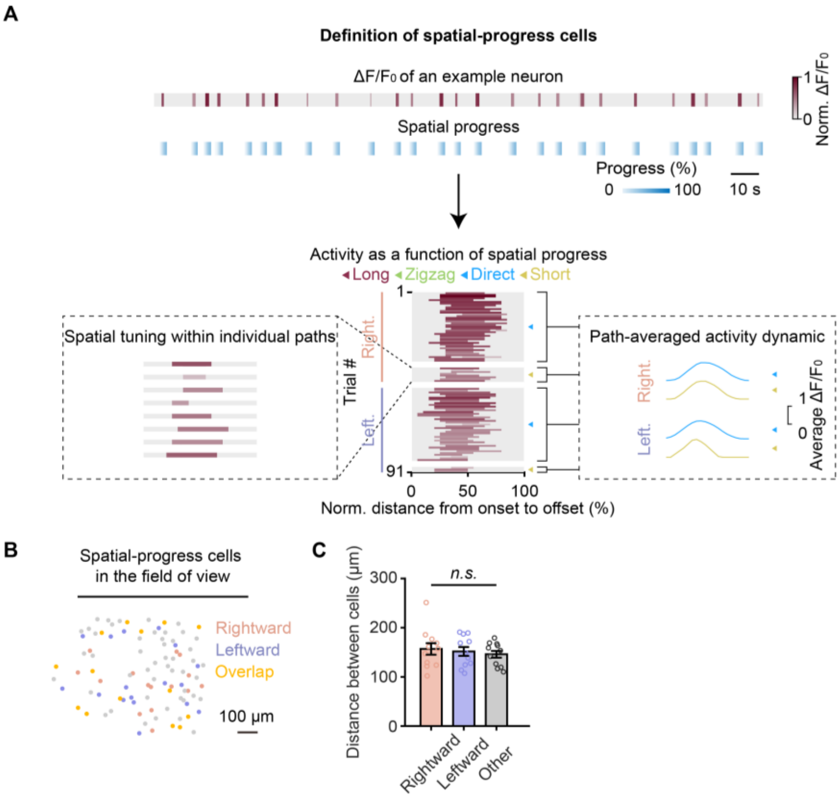


**Figure S5. Selection criteria for spatial-progress cells. Related to Figure 2.**

(A) Definition of spatial-progress cells. Top, activity of an example neuron and a raster plot showing the relative spatial progress in each trial. Bottom, schematic illustrating the identification of spatial-progress cells by constructing activity maps aligned by normalized distance from onset to offset.

(B) Spatial distribution of spatial-progress cells in the field of view of an example animal. Light red, rightward spatial-progress cells; light blue, leftward spatial-progress cells; light orange, overlap of rightward and leftward spatial-progress cells.

(C) Comparison of pairwise spatial distances among rightward spatial-progress cells, leftward spatial-progress cells, and non-progress cells ($n$ = 11 mice).

*n.s.*, not significant.

Detailed statistics are presented in Table S2.

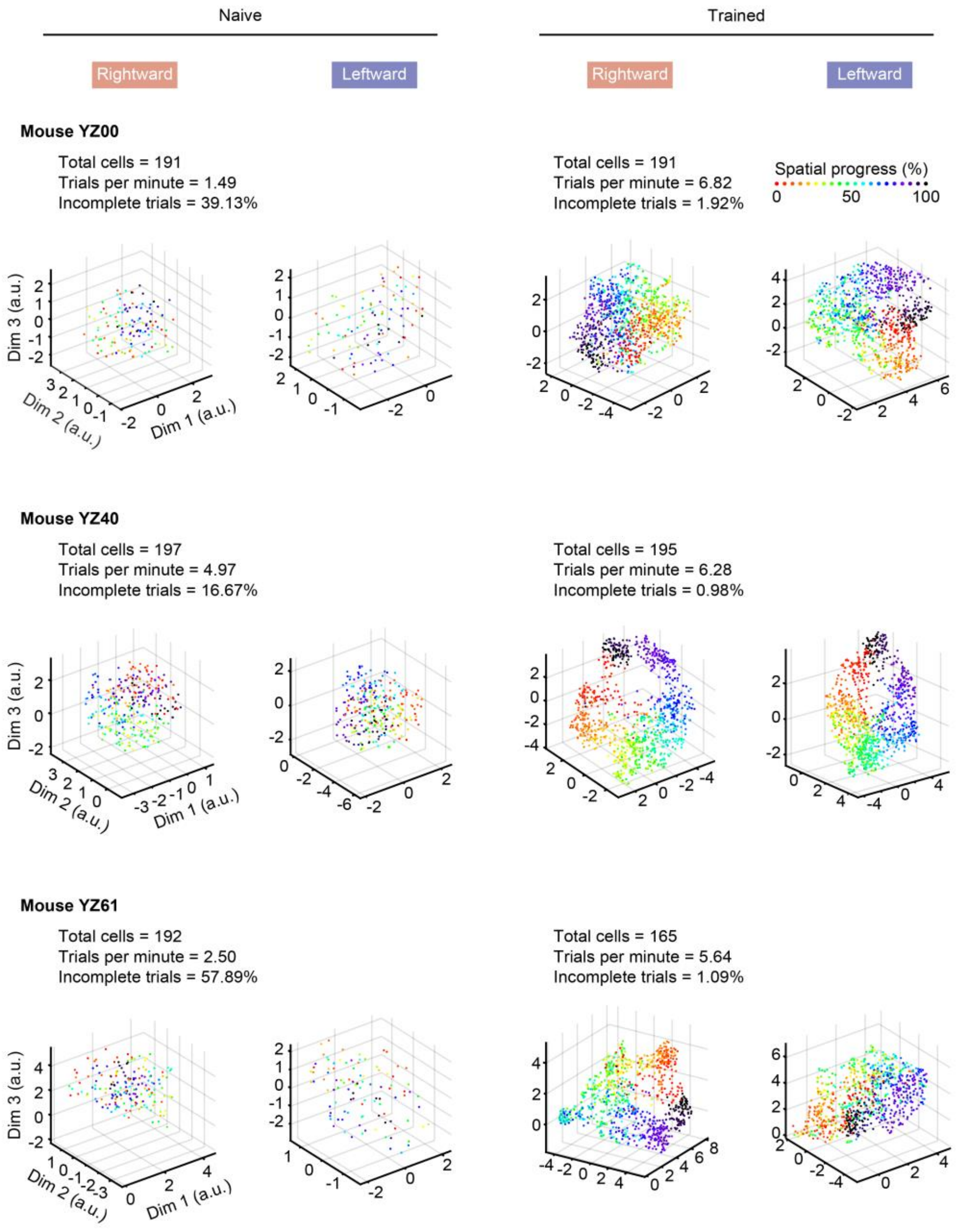


**Figure S6. Training reorganizes mPFC population activity in individual animals. Related to Figure 3.**

UMAP projections of population activity from example animals in the naive and trained stages. Manifolds were constructed separately from rightward or leftward trials. The total number of cells recorded, trials per minute, and proportion of incomplete trials are indicated. Rightward and leftward trials within the same animal showed remarkable consistency in the structure of manifolds, and similar sequential features were preserved across different animals. Thus, mPFC neuronal ensembles develop a robust sequential representation of spatial progress with learning.

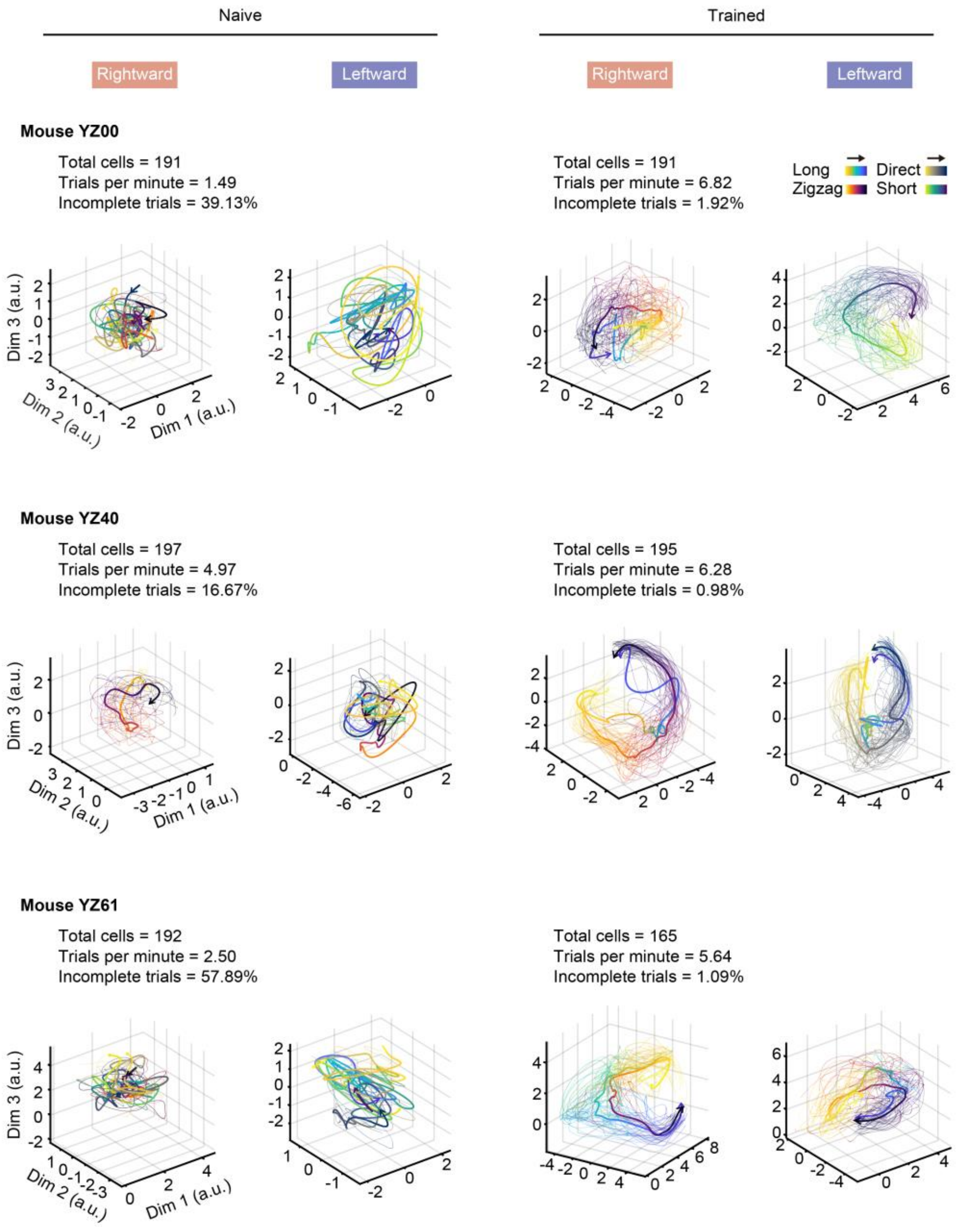


**Figure S7. Neural trajectories from different paths exhibit a similar ring-shaped characteristic. Related to Figure 3.**

Neural trajectories on the manifolds shown in Figure S6. Neural trajectories are grouped by paths and color-coded by both path type and spatial progress. Thick arrows, direction of the mean trajectory for each path along spatial progress. Fine lines, neural trajectories of individual trials. In the trained stage, trajectories from different paths exhibited a similar ring-shaped structure. The same progress stages were spatially close across paths, revealing a ring-shaped organization of population activity along spatial progress irrespective of path geometry. This structured organization was not observed in the naive stage.

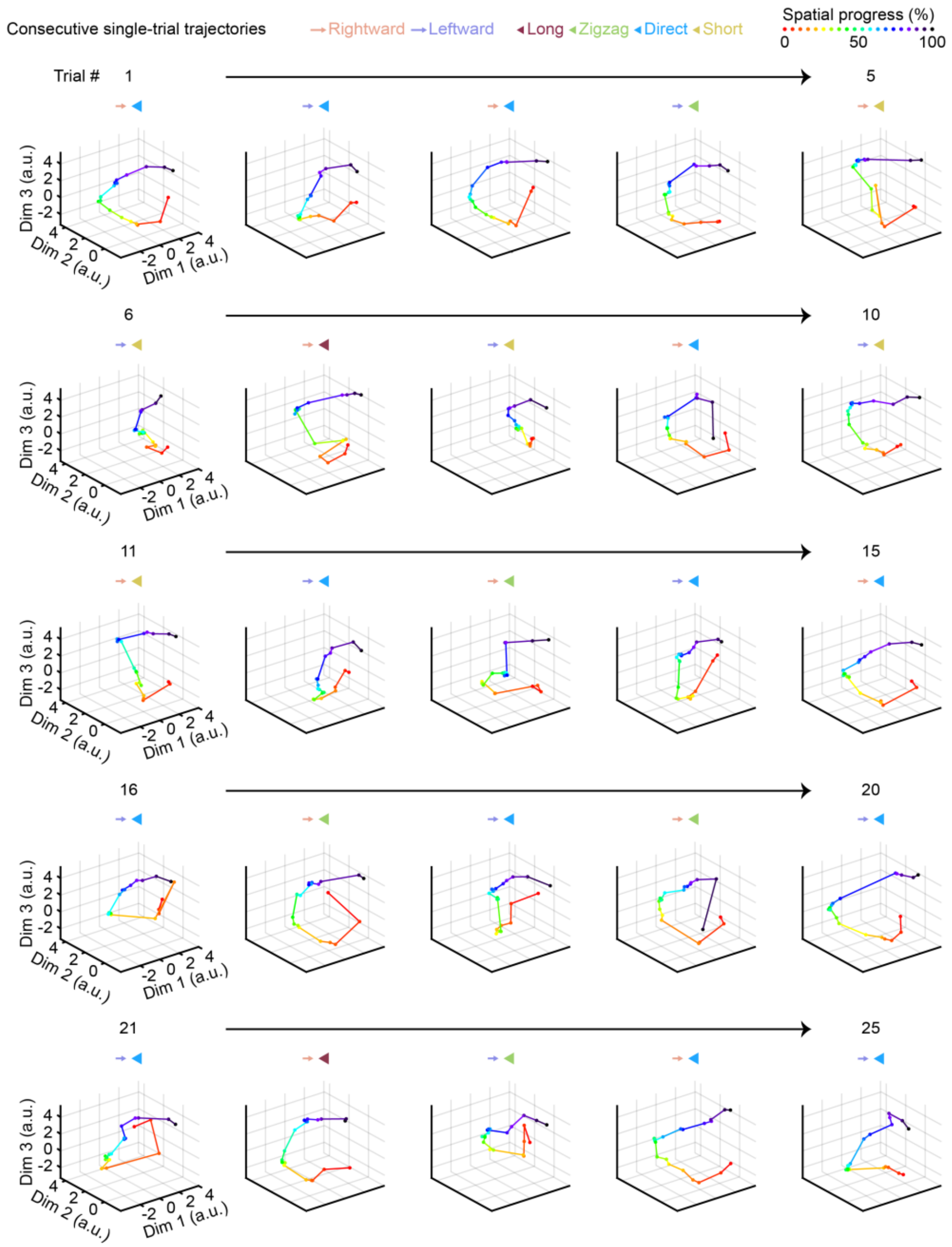


**Figure S8. Similar single-trial neural trajectories across directions and paths. Related to Figure 3.**

Consecutive single-trial neural trajectories from an example animal visualized on the UMAP manifold, colored by spatial progress. Neural trajectories were constructed from single trials of different running directions and visiting paths. All of the neural trajectories exhibited a similar sequential ring-shaped characteristic in the low-dimensional space (Video S2). Thus, mPFC neural ensembles encode the spatial progress from start to goal, independent of path geometry.

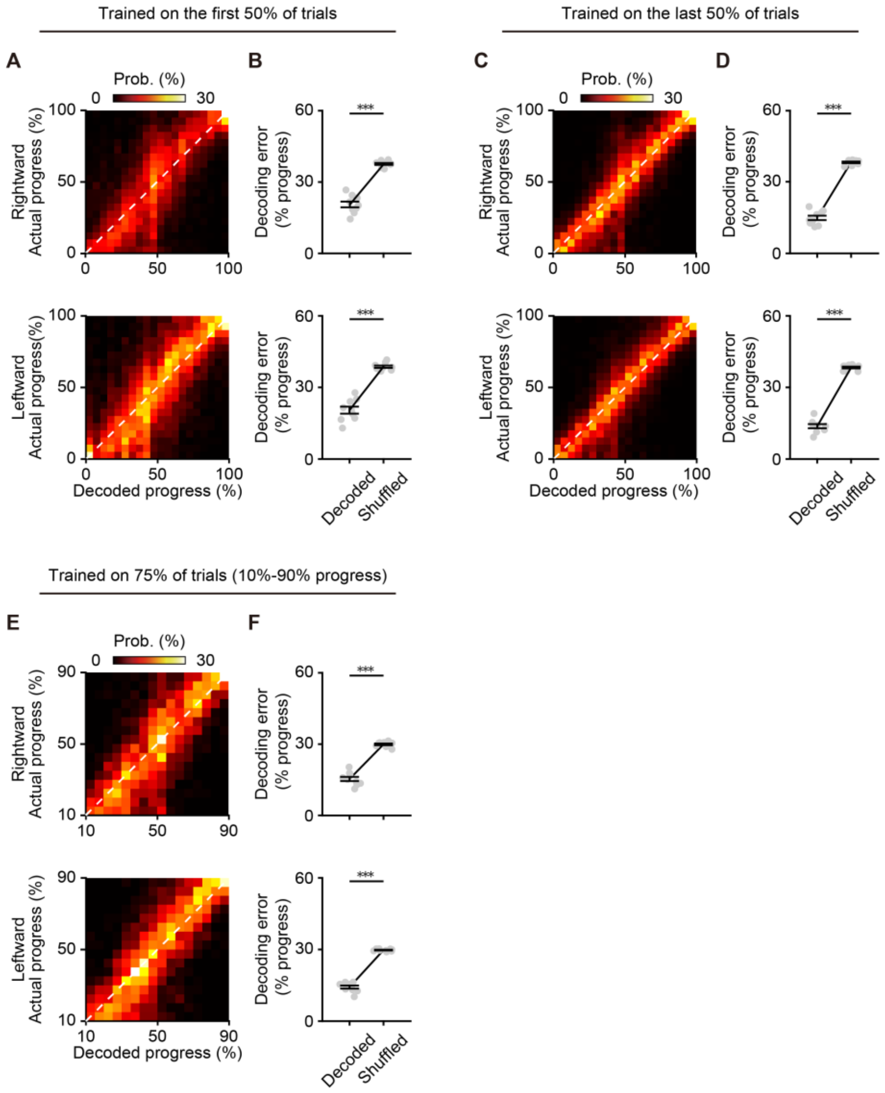


**Figure S9. Decoding for spatial progress is stable across the whole recording session. Related to Figure 3.**

(A) Confusion matrices showing decoded spatial progress, with the decoder trained on the first half of trials and tested on the second half. Top, rightward trials; bottom, leftward trials.
(B) Comparison of the decoding error between actual and shuffled datasets from (A) (*n* = 9 mice).
(C) Confusion matrices showing decoded spatial progress, with the decoder trained on the second half of trials and tested on the first half. Top, rightward trials; bottom, leftward trials.
(D) Comparison of the decoding error between actual and shuffled datasets from (C) (*n* = 9 mice).
(E) Confusion matrices showing decoded spatial progress using the dataset of 10%–90% progress. Top, rightward trials; bottom, leftward trials.
(F) Comparison of the decoding error between actual and shuffled datasets from (E) (*n* = 9 mice).
***$P$<0.001. Detailed statistics are presented in Table S2.

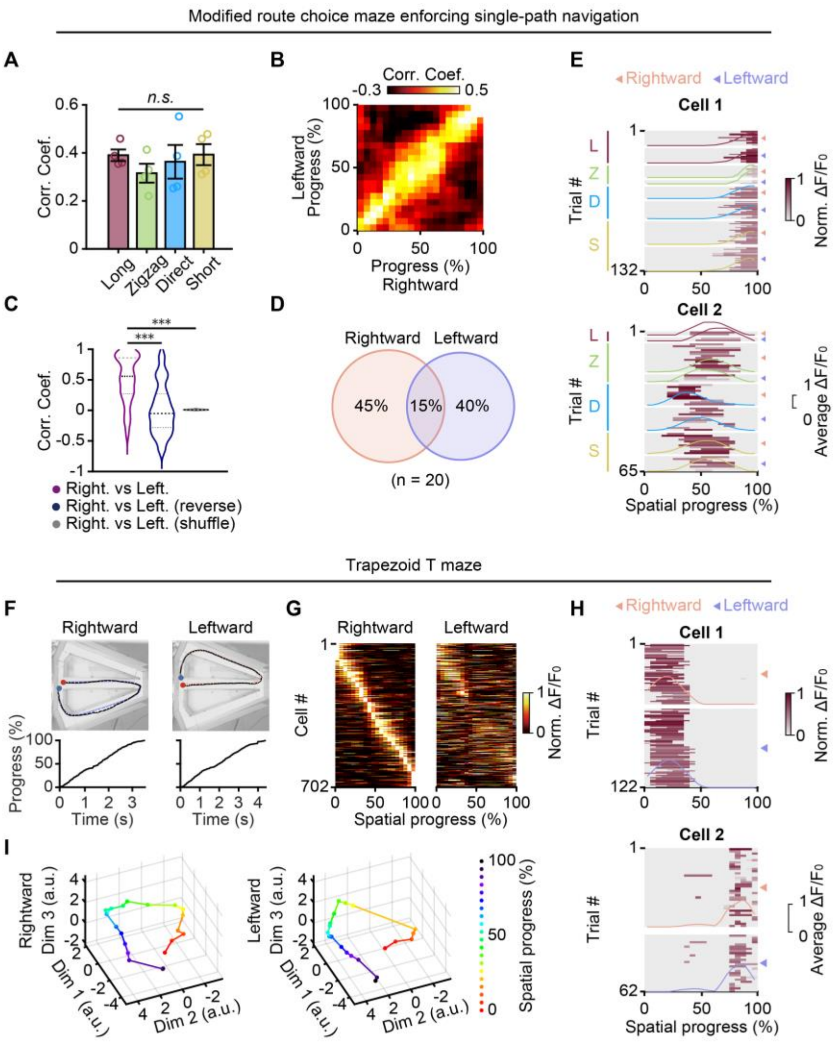


**Figure S10. Representation of spatial progress generalizes across contexts. Related to Figure 4.**

(A) Average correlation coefficients of normalized peak positions across trials within individual paths ($n$ = 4 mice).

(B) Population correlation matrix of trial-averaged activity between rightward and leftward trials.

(C) Comparison of the correlation coefficients between rightward and leftward trials with those after reversing or shuffling neuronal activity in leftward trials ($n$ = 128 cells).

(D) Venn diagram illustrating the relationship of spatial-progress cells between rightward and leftward trials.

(E) Activity maps of two example neurons as a function of spatial progress in the no-choice task (format as in Figure 4F). (A) to (E) are from the modified route choice paradigm enforcing single-path navigation, with the open path alternated per session.

(F) Example movement trajectories from each path (top) and the corresponding spatial progress as a function of elapsed time (bottom). Colored dots indicate trial onset (red) and trial offset (blue).

(G) Heatmaps showing trial-averaged neuronal activity aligned by spatial progress. Left, rightward trials; right, leftward trials. Neurons for both rightward and leftward trials are sorted by peak firing position in rightward trials. Only neurons active in both paths are shown.

(H) Activity maps of two example neurons as a function of spatial progress in the trapezoid T-maze (format as in Figure 4L).

(I) Single-trial neural trajectories from an example animal visualized on the UMAP manifold, colored by spatial progress. Similar to the route choice paradigm (Figure S8), neural trajectories from trials of different paths also exhibited a similar sequential ring-like feature. (F) to (I) are from the trapezoid T-maze.

***$P<0.001$. *n.s.*, not significant.

Detailed statistics are presented in Table S2.

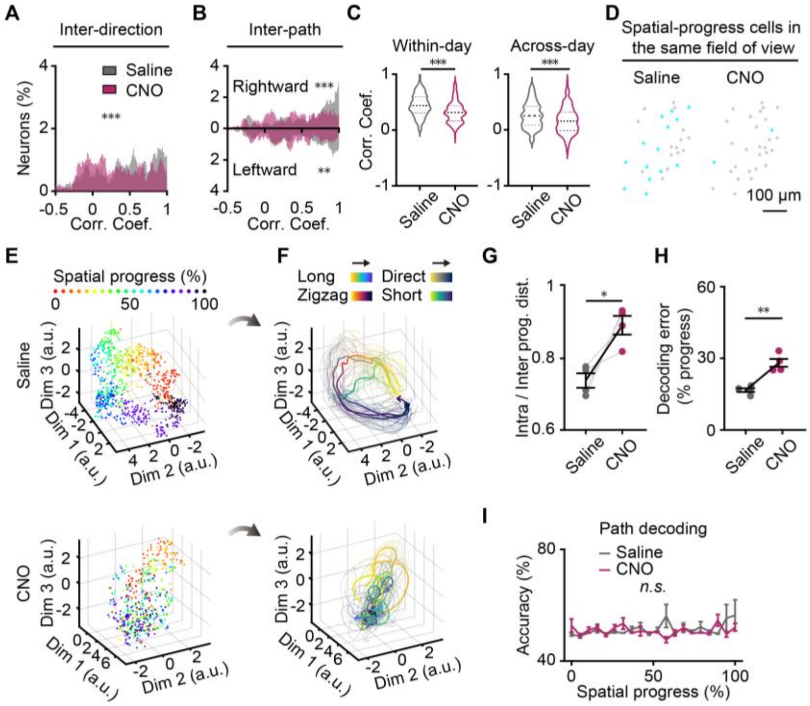


**Figure S11. Inactivation of CA1→mPFC disrupts the navigational efficiency and progress coding of mPFC neurons across environments. Related to Figures 5 and 6.**

(A, B) Distributions of the correlation coefficients between neuronal activity averaged across trials of different directions (A) or paths (B) (n = 223 cells).

(C) Comparison of correlation coefficients for the same path type before and after inactivation of the CA1→mPFC projection ($n$ = 223 cells). Left, within days; right, across days.

(D) Spatial distributions of spatial-progress cells from an example animal before (left) and after (right) inactivation of the CA1→mPFC projection.

(E–I) Same as Figures 6G–K, but for leftward trials ($n$ = 4 mice).

*$P$<0.05, **$P$<0.01 and ***$P$<0.001. *n.s.*, not significant.

Detailed statistics are presented in Table S2.
