## Supplementary Tables for "A Generalized Spatial Progress Code for Navigation in the Medial Prefrontal Cortex"

### Table S1. Extended statistical information for Figures 1 to 6.

| Figure panel | n/group | Primary statistic | *Post-hoc* test | Comparison | p value | Statistic |
|---|---|---|---|---|---|---|
| **Figure 1B** | n = 11 mice | Two-tailed paired Student's t-test | | Naive vs. Trained | <0.0001 | $t_{10}$ = -7.8622 |
| **Figure 1C** | n = 11 mice | Two-tailed paired Student's t-test | | Naive vs. Trained | <0.0001 | $t_{10}$ = 11.2180 |
| **Figure 1D** | n = 11 mice | Two-tailed paired Student's t-test | | Naive vs. Trained | 0.0002 | $t_{10}$ = 5.7145 |
| **Figure 1E** | n = 11 mice | Two-tailed paired Student's t-test | | Naive vs. Trained | 0.0002 | $t_{10}$ = 5.8241 |
| **Figure 1G** | n = 11 mice (naive) | One-way RM ANOVA | Bonferroni's multiple comparison | Main effect of group | 0.9391 | $F_{(3,30)}$ = 0.1339 |
| | | | | Long vs. Zigzag | >0.9999 | |
| | | | | Long vs. Direct | >0.9999 | |
| | | | | Long vs. Short | >0.9999 | |
| | | | | Zigzag vs. Direct | >0.9999 | |
| | | | | Zigzag vs. Short | >0.9999 | |
| | | | | Direct vs. Short | >0.9999 | |
| | n = 11 mice (trained) | One-way RM ANOVA | Bonferroni's multiple comparison | Main effect of group | 0.0002 | $F_{(3,30)}$ = 9.0539 |
| | | | | Long vs. Zigzag | 0.1465 | |
| | | | | Long vs. Direct | 0.0004 | |
| | | | | Long vs. Short | 0.0619 | |
| | | | | Zigzag vs. Direct | 0.0412 | |
| | | | | Zigzag vs. Short | >0.9999 | |
| | | | | Direct vs. Short | 0.5487 | |
| **Figure 1I** | n = 11 mice | Two-tailed paired Student's t-test | | Naive vs. Trained | 0.0011 | $t_{10}$ = -4.5278 |
| **Figure 1K** | Saline, n = 8 mice CNO, n = 8 mice | Two-way RM ANOVA | Bonferroni's multiple comparison | Group × Stage interaction | 0.0525 | $F_{(1,14)}$ = 0.4895 |
| | | | | Saline vs. CNO (naive) | 0.5578 | |
| | | | | Saline vs. CNO (trained) | 0.0048 | |
| **Figure 1L** | Saline, n = 8 mice CNO, n = 8 mice | Two-way RM ANOVA | Bonferroni's multiple comparison | Group × Stage interaction | 0.1725 | $F_{(1,14)}$ = 2.0673 |
| | | | | Saline vs. CNO (naive) | 0.4949 | |
| | | | | Saline vs. CNO (trained) | 0.0462 | |
| **Figure 1M** | Saline, n = 8 mice CNO, n = 8 mice | Two-way RM ANOVA | Bonferroni's multiple comparison | Group × Stage interaction | 0.2582 | $F_{(1,14)}$ = 1.3911 |
| | | | | Saline vs. CNO (naive) | 0.9175 | |
| | | | | Saline vs. CNO (trained) | <0.0001 | |
| **Figure 1N** | Saline, n = 8 mice CNO, n = 8 mice | Two-way RM ANOVA | Bonferroni's multiple comparison | Group × Stage interaction | 0.3425 | $F_{(1,14)}$ = 0.9652 |
| | | | | Saline vs. CNO (naive) | 0.5223 | |
| | | | | Saline vs. CNO (trained) | 0.3413 | |
| **Figure 1P** | Saline, n = 8 mice CNO, n = 8 mice | Two-way RM ANOVA | Bonferroni's multiple comparison | Group × Stage interaction | 0.6015 | $F_{(1,14)}$ = 0.2856 |
| | | | | Saline vs. CNO (naive) | 0.3257 | |
| | | | | Saline vs. CNO (trained) | 0.9076 | |
| **Figure 2E** | n = 1023 cells (naive) and n = 891 cells (trained) from 11 mice | Two-way RM ANOVA | Bonferroni's multiple comparison | Stage × Alignment condition interaction | <0.0001 | $F_{(2,1788)}$ = 26.4210 |
| | | | | Time vs. Distance (naive) | >0.9999 | |
| | | | | Time vs. Progress (naive) | 0.1132 | |
| | | | | Distance vs. Progress (naive) | 0.0538 | |
| | | | | Time vs. Distance (trained) | <0.0001 | |
| | | | | Time vs. Progress (trained) | <0.0001 | |
| | | | | Distance vs. Progress (trained) | <0.0001 | |
| **Figure 2G** | n = 255 spatial-progress cells (trained) from 11 mice | One-way RM ANOVA | Bonferroni's multiple comparison | Main effect of group | <0.0001 | $F_{(2,508)}$ = 26.2899 |
| | | | | Time vs. Distance | 0.7220 | |
| | | | | Time vs. Progress | <0.0001 | |
| | | | | Distance vs. Progress | <0.0001 | |
| **Figure 2H** | n = 11 mice (trained) | Two-tailed paired Student's t-test | | Naive vs. Trained (rightward) | 0.0005 | $t_{10}$ = -5.0262 |
| | | | | Naive vs. Trained (leftward) | 0.0006 | $t_{10}$ = -4.9542 |
| **Figure 2K** | n = 64 spatial-progress cells (naive) and n = 255 spatial-progress cells (trained) from 11 mice | Two-sample K-S test | | Naive vs. Trained | <0.0001 | D = 0.1921 |
| **Figure 2L** | n = 64 spatial-progress cells (naive) and n = 255 spatial-progress cells (trained) from 11 mice | Mann-Whitney U test | | Naive vs. Trained | <0.0001 | U = 5632 |
| **Figure 2N** | n = 11 mice | Two-way RM ANOVA | Bonferroni's multiple comparison | Stage × Condition interaction | 0.0217 | $F_{(1,10)}$ = 7.3801 |
| | | | | Real vs. Shuffle (naive) | >0.9999 | |
| | | | | Real vs. Shuffle (trained) | 0.0132 | |
| | | | | Naive vs. Trained (real) | 0.0002 | |

| Figure panel | n/group | Primary statistic | *Post-hoc* test | Comparison | p value | Statistic |
|---|---|---|---|---|---|---|
| **Figure 3E** | n = 9 mice (trained) | Two-way RM ANOVA | Bonferroni's multiple comparison | Direction × Distance interaction | 0.8405 | $F_{(1,16)}$ = 0.0419 |
| | | | | Intra- vs. Inter-bin distance (rightward) | <0.0001 | |
| | | | | Intra- vs. Inter-bin distance (leftward) | <0.0001 | |
| **Figure 3F** | n = 9 mice (trained) | Two-way RM ANOVA | Bonferroni's multiple comparison | Direction × Distance interaction | 0.6642 | $F_{(1,16)}$ = 0.1957 |
| | | | | Intra- vs. Inter-path distance (rightward) | 0.2302 | |
| | | | | Intra- vs. Inter-path distance (leftward) | 0.2592 | |
| **Figure 3G** | n = 9 mice (trained) | Two-tailed paired Student's t-test | | Intra- vs. Inter-direction distance | 0.0130 | $t_8$ = -3.1788 |
| **Figure 3I** | n = 9 mice (trained) | Two-tailed paired Student's t-test | | Decoded vs. Shuffled error (rightward) | <0.0001 | $t_8$ = -15.2622 |
| | | | | Decoded vs. Shuffled error (leftward) | <0.0001 | $t_8$ = -17.5144 |
| **Figure 3K** | n = 9 mice (trained) | Two-tailed paired Student's t-test | | Decoded vs. Shuffled error (rightward) | <0.0001 | $t_8$ = -9.5564 |
| | | | | Decoded vs. Shuffled error (leftward) | <0.0001 | $t_8$ = -9.5847 |
| **Figure 3L (top)** | n = 9 mice (trained) | Two-way RM ANOVA | Bonferroni's multiple comparison | Group × Progress stage interaction | 0.3250 | $F_{(19,152)}$ = 1.1320 |
| | | | | Decoded vs. Shuffled accuracy (0% – 5%) | >0.9999 | |
| | | | | Decoded vs. Shuffled accuracy (5% – 10%) | >0.9999 | |
| | | | | Decoded vs. Shuffled accuracy (10% – 15%) | >0.9999 | |
| | | | | Decoded vs. Shuffled accuracy (15% – 20%) | 0.2243 | |
| | | | | Decoded vs. Shuffled accuracy (20% – 25%) | >0.9999 | |
| | | | | Decoded vs. Shuffled accuracy (25% – 30%) | 0.9782 | |
| | | | | Decoded vs. Shuffled accuracy (30% – 35%) | 0.0093 | |
| | | | | Decoded vs. Shuffled accuracy (35% – 40%) | >0.9999 | |
| | | | | Decoded vs. Shuffled accuracy (40% – 45%) | >0.9999 | |
| | | | | Decoded vs. Shuffled accuracy (45% – 50%) | >0.9999 | |
| | | | | Decoded vs. Shuffled accuracy (50% – 55%) | >0.9999 | |
| | | | | Decoded vs. Shuffled accuracy (55% – 60%) | 0.8907 | |
| | | | | Decoded vs. Shuffled accuracy (60% – 65%) | >0.9999 | |
| | | | | Decoded vs. Shuffled accuracy (65% – 70%) | >0.9999 | |
| | | | | Decoded vs. Shuffled accuracy (70% – 75%) | >0.9999 | |
| | | | | Decoded vs. Shuffled accuracy (75% – 80%) | >0.9999 | |
| | | | | Decoded vs. Shuffled accuracy (80% – 85%) | >0.9999 | |
| | | | | Decoded vs. Shuffled accuracy (85% – 90%) | >0.9999 | |
| | | | | Decoded vs. Shuffled accuracy (90% – 95%) | >0.9999 | |
| | | | | Decoded vs. Shuffled accuracy (95% – 100%) | >0.9999 | |
| **Figure 3L (bottom)** | n = 9 mice (trained) | Two-way RM ANOVA | Bonferroni's multiple comparison | Group × Progress stage interaction | 0.4997 | $F_{(19,152)}$ = 0.9697 |
| | | | | Decoded vs. Shuffled accuracy (0% – 5%) | >0.9999 | |
| | | | | Decoded vs. Shuffled accuracy (5% – 10%) | >0.9999 | |
| | | | | Decoded vs. Shuffled accuracy (10% – 15%) | >0.9999 | |
| | | | | Decoded vs. Shuffled accuracy (15% – 20%) | 0.2290 | |
| | | | | Decoded vs. Shuffled accuracy (20% – 25%) | >0.9999 | |
| | | | | Decoded vs. Shuffled accuracy (25% – 30%) | >0.9999 | |
| | | | | Decoded vs. Shuffled accuracy (30% – 35%) | >0.9999 | |
| | | | | Decoded vs. Shuffled accuracy (35% – 40%) | >0.9999 | |
| | | | | Decoded vs. Shuffled accuracy (40% – 45%) | 0.9528 | |
| | | | | Decoded vs. Shuffled accuracy (45% – 50%) | >0.9999 | |
| | | | | Decoded vs. Shuffled accuracy (50% – 55%) | >0.9999 | |
| | | | | Decoded vs. Shuffled accuracy (55% – 60%) | >0.9999 | |
| | | | | Decoded vs. Shuffled accuracy (60% – 65%) | >0.9999 | |
| | | | | Decoded vs. Shuffled accuracy (65% – 70%) | >0.9999 | |
| | | | | Decoded vs. Shuffled accuracy (70% – 75%) | >0.9999 | |
| | | | | Decoded vs. Shuffled accuracy (75% – 80%) | >0.9999 | |
| | | | | Decoded vs. Shuffled accuracy (80% – 85%) | 0.7198 | |
| | | | | Decoded vs. Shuffled accuracy (85% – 90%) | 0.0419 | |
| | | | | Decoded vs. Shuffled accuracy (90% – 95%) | >0.9999 | |
| | | | | Decoded vs. Shuffled accuracy (95% – 100%) | >0.9999 | |

| Figure panel | n/group | Primary statistic | *Post-hoc* test | Comparison | p value | Statistic |
|---|---|---|---|---|---|---|
| **Figure 3M** | n = 9 mice (trained) | Two-way RM ANOVA | Bonferroni's multiple comparison | Group × Progress stage interaction | <0.0001 | $F_{(19,152)}$ = 3.9466 |
| | | | | Decoded vs. Shuffled accuracy (0% – 5%) | >0.9999 | |
| | | | | Decoded vs. Shuffled accuracy (5% – 10%) | 0.0982 | |
| | | | | Decoded vs. Shuffled accuracy (10% – 15%) | >0.9999 | |
| | | | | Decoded vs. Shuffled accuracy (15% – 20%) | 0.4927 | |
| | | | | Decoded vs. Shuffled accuracy (20% – 25%) | 0.0448 | |
| | | | | Decoded vs. Shuffled accuracy (25% – 30%) | >0.9999 | |
| | | | | Decoded vs. Shuffled accuracy (30% – 35%) | >0.9999 | |
| | | | | Decoded vs. Shuffled accuracy (35% – 40%) | 0.3986 | |
| | | | | Decoded vs. Shuffled accuracy (40% – 45%) | 0.2734 | |
| | | | | Decoded vs. Shuffled accuracy (45% – 50%) | 0.0904 | |
| | | | | Decoded vs. Shuffled accuracy (50% – 55%) | 0.5199 | |
| | | | | Decoded vs. Shuffled accuracy (55% – 60%) | 0.0035 | |
| | | | | Decoded vs. Shuffled accuracy (60% – 65%) | 0.1963 | |
| | | | | Decoded vs. Shuffled accuracy (65% – 70%) | 0.2955 | |
| | | | | Decoded vs. Shuffled accuracy (70% – 75%) | 0.0719 | |
| | | | | Decoded vs. Shuffled accuracy (75% – 80%) | 0.0027 | |
| | | | | Decoded vs. Shuffled accuracy (80% – 85%) | 0.0008 | |
| | | | | Decoded vs. Shuffled accuracy (85% – 90%) | <0.0001 | |
| | | | | Decoded vs. Shuffled accuracy (90% – 95%) | <0.0001 | |
| | | | | Decoded vs. Shuffled accuracy (95% – 100%) | <0.0001 | |
| **Figure 4C** | n = 4 mice | Two-tailed paired Student’s t-test | | Real vs. Shuffle (long–zigzag) | 0.0300 | $t_3$ = 3.8957 |
| | | | | Real vs. Shuffle (long–direct) | 0.0032 | $t_3$ = 8.7050 |
| | | | | Real vs. Shuffle (long–short) | 0.0057 | $t_3$ = 7.1257 |
| | | | | Real vs. Shuffle (zigzag–direct) | 0.0064 | $t_3$ = 6.8354 |
| | | | | Real vs. Shuffle (zigzag–short) | 0.0042 | $t_3$ = 7.9310 |
| | | | | Real vs. Shuffle (direct–short) | 0.0064 | $t_3$ = 6.8329 |
| **Figure 4D** | n = 128 cells from 4 mice | Wilcoxon signed rank test | | Real vs. Shuffle (long–zigzag) | <0.0001 | Z = 6.7214 |
| | | | | Real vs. Shuffle (long–direct) | <0.0001 | Z = 6.1720 |
| | | | | Real vs. Shuffle (long–short) | <0.0001 | Z = 6.9585 |
| | | | | Real vs. Shuffle (zigzag–direct) | <0.0001 | Z = 6.2641 |
| | | | | Real vs. Shuffle (zigzag–short) | <0.0001 | Z = 7.0489 |
| | | | | Real vs. Shuffle (direct–short) | <0.0001 | Z = 5.8245 |
| **Figure 4G** | n = 4 mice | Two-tailed paired Student’s t-test | | Decoded vs. Shuffled error (test set: long) | 0.0215 | $t_3$ = -4.4198 |
| | | | | Decoded vs. Shuffled error (test set: zigzag) | 0.0547 | $t_3$ = -3.0661 |
| | | | | Decoded vs. Shuffled error (test set: direct) | 0.0359 | $t_3$ = -3.6343 |
| | | | | Decoded vs. Shuffled error (test set: short) | 0.0304 | $t_3$ = -3.8760 |
| **Figure 4J** | n = 771 cells from 8 mice | Wilcoxon signed rank test | | Left–Right vs. Left–Right (shuffle) | <0.0001 | Z = 18.5026 |
| **Figure 4O** | n = 8 mice | Two-tailed paired Student’s t-test | | Decoded vs. Shuffled error | 0.0002 | $t_7$ = -7.3204 |
| **Figure 4P** | n = 8 mice | Two-way RM ANOVA | Bonferroni's multiple comparison | Group × Progress stage interaction | 0.4332 | $F_{(19,133)}$ = 1.0287 |
| | | | | Decoded vs. Shuffled accuracy (0% – 5%) | >0.9999 | |
| | | | | Decoded vs. Shuffled accuracy (5% – 10%) | >0.9999 | |
| | | | | Decoded vs. Shuffled accuracy (10% – 15%) | >0.9999 | |
| | | | | Decoded vs. Shuffled accuracy (15% – 20%) | >0.9999 | |
| | | | | Decoded vs. Shuffled accuracy (20% – 25%) | >0.9999 | |
| | | | | Decoded vs. Shuffled accuracy (25% – 30%) | >0.9999 | |
| | | | | Decoded vs. Shuffled accuracy (30% – 35%) | >0.9999 | |
| | | | | Decoded vs. Shuffled accuracy (35% – 40%) | >0.9999 | |
| | | | | Decoded vs. Shuffled accuracy (40% – 45%) | >0.9999 | |
| | | | | Decoded vs. Shuffled accuracy (45% – 50%) | >0.9999 | |
| | | | | Decoded vs. Shuffled accuracy (50% – 55%) | >0.9999 | |
| | | | | Decoded vs. Shuffled accuracy (55% – 60%) | >0.9999 | |
| | | | | Decoded vs. Shuffled accuracy (60% – 65%) | 0.5501 | |
| | | | | Decoded vs. Shuffled accuracy (65% – 70%) | 0.7068 | |
| | | | | Decoded vs. Shuffled accuracy (70% – 75%) | 0.2148 | |
| | | | | Decoded vs. Shuffled accuracy (75% – 80%) | >0.9999 | |
| | | | | Decoded vs. Shuffled accuracy (80% – 85%) | 0.3392 | |
| | | | | Decoded vs. Shuffled accuracy (85% – 90%) | 0.7536 | |
| | | | | Decoded vs. Shuffled accuracy (90% – 95%) | >0.9999 | |
| | | | | Decoded vs. Shuffled accuracy (95% – 100%) | >0.9999 | |
| **Figure 4S** | n = 8 mice | Two-way RM ANOVA | Bonferroni's multiple comparison | Direction × Distance interaction | 0.4359 | $F_{(1,7)}$ = 0.6828 |
| | | | | Intra- vs. Inter-bin distance (rightward) | 0.0006 | |
| | | | | Intra- vs. Inter-bin distance (leftward) | 0.0017 | |
| **Figure 4T** | n = 8 mice | Two-tailed paired Student’s t-test | | Intra- vs. Inter-path distance | 0.0869 | $t_7$ = -1.9902 |
| **Figure 5C** | n = 4 mice | Two-tailed paired Student’s t-test | | Saline vs. CNO | 0.0190 | $t_3$ = 4.6267 |
| **Figure 5D** | n = 4 mice | Two-tailed paired Student’s t-test | | Saline vs. CNO | 0.0301 | $t_3$ = -3.8911 |

| Figure panel | n/group | Primary statistic | *Post-hoc* test | Comparison | p value | Statistic |
|---|---|---|---|---|---|---|
| **Figure 5E** | n = 4 mice | Two-tailed paired Student's t-test | | Saline vs. CNO | 0.0158 | $t_3$ = -4.9535 |
| **Figure 5G** | n = 4 mice | Two-tailed paired Student's t-test | | Saline vs. CNO | 0.2925 | $t_3$ = 1.2736 |
| **Figure 5H** | n = 4 mice | Two-tailed paired Student's t-test | | Saline vs. CNO | 0.1236 | $t_3$ = -2.1251 |
| **Figure 5I** | n = 4 mice | Two-tailed paired Student's t-test | | Saline vs. CNO | 0.8448 | $t_3$ = 0.2132 |
| **Figure 6C** | n = 59 spatial-progress cells (identified in either saline or CNO stage) from 4 mice | Wilcoxon signed rank test | | Saline vs. CNO | 0.0007 | $t_{58}$ = -3.5781 |
| **Figure 6F** | n = 63 spatial-progress cells (saline) and n = 31 spatial-progress cells (CNO) from 4 mice | Two-sample K-S test | | Saline vs. CNO (field distribution) | 0.5592 | D = 0.0919 |
| | | Mann-Whitney U test | | Saline vs. CNO (field width) | 0.2126 | U = 826 |
| **Figure 6I** | n = 4 mice | Two-tailed paired Student's t-test | | Saline vs. CNO (rightward) | 0.0240 | $t_3$ = -4.2420 |
| **Figure 6J** | n = 4 mice | Two-tailed paired Student's t-test | | Saline vs. CNO (rightward) | 0.0142 | $t_3$ = -5.1448 |
| **Figure 6K** | n = 4 mice | Two-way RM ANOVA | Bonferroni's multiple comparison | Treatment × Progress stage interaction | 0.4497 | $F_{(19,57)}$ = 1.0242 |
| | | | | Accuracy of Saline vs. CNO (0% – 5%) | >0.9999 | |
| | | | | Accuracy of Saline vs. CNO (5% – 10%) | >0.9999 | |
| | | | | Accuracy of Saline vs. CNO (10% – 15%) | >0.9999 | |
| | | | | Accuracy of Saline vs. CNO (15% – 20%) | >0.9999 | |
| | | | | Accuracy of Saline vs. CNO (20% – 25%) | >0.9999 | |
| | | | | Accuracy of Saline vs. CNO (25% – 30%) | >0.9999 | |
| | | | | Accuracy of Saline vs. CNO (30% – 35%) | >0.9999 | |
| | | | | Accuracy of Saline vs. CNO (35% – 40%) | >0.9999 | |
| | | | | Accuracy of Saline vs. CNO (40% – 45%) | >0.9999 | |
| | | | | Accuracy of Saline vs. CNO (45% – 50%) | >0.9999 | |
| | | | | Accuracy of Saline vs. CNO (50% – 55%) | >0.9999 | |
| | | | | Accuracy of Saline vs. CNO (55% – 60%) | >0.9999 | |
| | | | | Accuracy of Saline vs. CNO (60% – 65%) | >0.9999 | |
| | | | | Accuracy of Saline vs. CNO (65% – 70%) | 0.0169 | |
| | | | | Accuracy of Saline vs. CNO (70% – 75%) | >0.9999 | |
| | | | | Accuracy of Saline vs. CNO (75% – 80%) | >0.9999 | |
| | | | | Accuracy of Saline vs. CNO (80% – 85%) | >0.9999 | |
| | | | | Accuracy of Saline vs. CNO (85% – 90%) | >0.9999 | |
| | | | | Accuracy of Saline vs. CNO (90% – 95%) | >0.9999 | |
| | | | | Accuracy of Saline vs. CNO (95% – 100%) | >0.9999 | |

### Table S2. Extended statistical information for Figures S1 to S11.

| Figure panel | n/group | Primary statistic | *Post-hoc* test | Comparison | p value | Statistic |
|---|---|---|---|---|---|---|
| **Figure S1E** | n = 11 mice | Two-way RM ANOVA | Bonferroni's multiple comparison | Path × Stage interaction | 0.2177 | $F_{(3,40)}$ = 1.5452 |
| | | | | Naive vs. Trained (long) | <0.0001 | |
| | | | | Naive vs. Trained (zigzag) | <0.0001 | |
| | | | | Naive vs. Trained (direct) | 0.0011 | |
| | | | | Naive vs. Trained (short) | 0.0195 | |
| **Figure S1F** | n = 11 mice | Two-way RM ANOVA | Bonferroni's multiple comparison | Direction × Stage interaction | 0.2315 | $F_{(1,20)}$ = 1.5226 |
| | | | | Naive vs. Trained (rightward) | <0.0001 | |
| | | | | Naive vs. Trained (leftward) | <0.0001 | |
| **Figure S1G** | n = 11 mice | Two-tailed paired Student's t-test | | Naive vs. Trained | 0.0059 | $t_{10}$ = 3.4848 |
| **Figure S1I** | n = 11 mice | Two-tailed paired Student's t-test | | Naive vs. Trained | <0.0001 | $t_{10}$ = 7.9719 |
| **Figure S1J** | Saline, n = 8 mice CNO, n = 8 mice | Two-way RM ANOVA | Bonferroni's multiple comparison | Group × Stage interaction | 0.9381 | $F_{(1,14)}$ = 0.0063 |
| | | | | Saline vs. CNO (naive) | 0.8463 | |
| | | | | Saline vs. CNO (trained) | 0.2824 | |
| **Figure S3C** | n = 1023 cells (naive) and n = 891 cells (trained) from 11 mice | Mann-Whitney U test | | Naive vs. Trained | <0.0001 | U = 292201 |
| **Figure S3D** | n = 1023 cells from 11 mice (naive) | One-way RM ANOVA | Bonferroni's multiple comparison | Main effect of group | <0.0001 | $F_{(2,1772)}$ = 216.5878 |
| | | | | Right–Left vs. Right–Left (reverse) | <0.0001 | |
| | | | | Right–Left vs. Right–Left (shuffle) | <0.0001 | |
| | n = 891 cells from 11 mice (trained) | One-way RM ANOVA | Bonferroni's multiple comparison | Main effect of group | <0.0001 | $F_{(2,1750)}$ = 307.8943 |
| | | | | Right–Left vs. Right–Left (reverse) | <0.0001 | |
| | | | | Right–Left vs. Right–Left (shuffle) | <0.0001 | |
| **Figure S3E** | n = 1023 cells (naive) and n = 891 cells (trained) from 11 mice | Mann-Whitney U test | | Naive vs. Trained (rightward) | <0.0001 | U = 164629 |
| | | | | Naive vs. Trained (leftward) | <0.0001 | U = 247285 |
| **Figure S3F** | n = 1023 cells (naive) and n = 891 cells (trained) from 11 mice | Mann-Whitney U test | | Naive vs. Trained (within-day) | <0.0001 | U = 176585 |
| | | | | Naive vs. Trained (across-day) | <0.0001 | U = 158018 |
| **Figure S5C** | n = 11 mice | One-way RM ANOVA | | Main effect of group | 0.4288 | $F_{(2,20)}$ = 0.8836 |
| | | | | Rightward vs. Leftward | >0.9999 | |
| | | | | Rightward vs. Other | 0.5968 | |
| | | | | Leftward vs. Other | >0.9999 | |
| **Figure S9B** | n = 9 mice (trained) | Two-tailed paired Student's t-test | | Decoded vs. Shuffled error (rightward) | <0.0001 | $t_8$ = -12.6197 |
| | | | | Decoded vs. Shuffled error (leftward) | <0.0001 | $t_8$ = -13.2841 |
| **Figure S9D** | n = 9 mice (trained) | Two-tailed paired Student's t-test | | Decoded vs. Shuffled error (rightward) | <0.0001 | $t_8$ = -17.7667 |
| | | | | Decoded vs. Shuffled error (leftward) | <0.0001 | $t_8$ = -20.5202 |
| **Figure S9F** | n = 9 mice (trained) | Two-tailed paired Student's t-test | | Decoded vs. Shuffled error (rightward) | <0.0001 | $t_8$ = -11.3364 |
| | | | | Decoded vs. Shuffled error (leftward) | <0.0001 | $t_8$ = -18.8576 |
| **Figure S10A** | n = 4 mice | One-way RM ANOVA | Bonferroni's multiple comparison | Main effect of group | 0.6508 | $F_{(3,12)}$ = 0.5612 |
| | | | | Long vs. Zigzag | >0.9999 | |
| | | | | Long vs. Direct | >0.9999 | |
| | | | | Long vs. Short | >0.9999 | |
| | | | | Zigzag vs. Direct | >0.9999 | |
| | | | | Zigzag vs. Short | >0.9999 | |
| | | | | Direct vs. Short | >0.9999 | |
| **Figure S10C** | n = 128 cells from 4 mice | One-way RM ANOVA | Bonferroni's multiple comparison | Main effect of group | <0.0001 | $F_{(2,254)}$ = 96.5207 |
| | | | | Right–Left vs. Right–Left (reverse) | <0.0001 | |
| | | | | Right–Left vs. Right–Left (shuffle) | <0.0001 | |
| **Figure S11A** | n = 223 cells from 4 mice | Wilcoxon signed rank test | | Saline vs. CNO | 0.0006 | Z = 3.4389 |
| **Figure S11B** | n = 223 cells from 4 mice | Wilcoxon signed rank test | | Saline vs. CNO (rightward) | 0.0001 | Z = -3.8184 |
| | | | | Saline vs. CNO (leftward) | 0.0011 | Z = -3.2564 |
| **Figure S11C** | n = 223 cells from 4 mice | Wilcoxon signed rank test | | Saline vs. CNO (within-day) | <0.0001 | Z = 5.9862 |
| | | | | Saline vs. CNO (across-day) | <0.0001 | Z = 4.1273 |
| **Figure S11G** | n = 4 mice | Two-tailed paired Student's t-test | | Saline vs. CNO (leftward) | 0.0385 | $t_3$ = -3.5362 |
| **Figure S11H** | n = 4 mice | Two-tailed paired Student's t-test | | Saline vs. CNO (leftward) | 0.0088 | $t_3$ = -6.1193 |

| Figure panel | n/group | Primary statistic | *Post-hoc* test | Comparison | p value | Statistic |
|---|---|---|---|---|---|---|
| **Figure S11I** | n = 4 mice | Two-way RM ANOVA | Bonferroni's multiple comparison | Treatment × Progress stage interaction | 0.2818 | $F_{(19,57)}$ = 1.2109 |
| | | | | Accuracy of Saline vs. CNO (0% – 5%) | >0.9999 | |
| | | | | Accuracy of Saline vs. CNO (5% – 10%) | >0.9999 | |
| | | | | Accuracy of Saline vs. CNO (10% – 15%) | >0.9999 | |
| | | | | Accuracy of Saline vs. CNO (15% – 20%) | >0.9999 | |
| | | | | Accuracy of Saline vs. CNO (20% – 25%) | >0.9999 | |
| | | | | Accuracy of Saline vs. CNO (25% – 30%) | >0.9999 | |
| | | | | Accuracy of Saline vs. CNO (30% – 35%) | >0.9999 | |
| | | | | Accuracy of Saline vs. CNO (35% – 40%) | >0.9999 | |
| | | | | Accuracy of Saline vs. CNO (40% – 45%) | >0.9999 | |
| | | | | Accuracy of Saline vs. CNO (45% – 50%) | >0.9999 | |
| | | | | Accuracy of Saline vs. CNO (50% – 55%) | >0.9999 | |
| | | | | Accuracy of Saline vs. CNO (55% – 60%) | 0.0681 | |
| | | | | Accuracy of Saline vs. CNO (60% – 65%) | >0.9999 | |
| | | | | Accuracy of Saline vs. CNO (65% – 70%) | >0.9999 | |
| | | | | Accuracy of Saline vs. CNO (70% – 75%) | >0.9999 | |
| | | | | Accuracy of Saline vs. CNO (75% – 80%) | >0.9999 | |
| | | | | Accuracy of Saline vs. CNO (80% – 85%) | >0.9999 | |
| | | | | Accuracy of Saline vs. CNO (85% – 90%) | >0.9999 | |
| | | | | Accuracy of Saline vs. CNO (90% – 95%) | 0.8431 | |
| | | | | Accuracy of Saline vs. CNO (95% – 100%) | >0.9999 | |
